# The deubiquitylase USP31 controls the Chromosomal Passenger Complex and spindle dynamics

**DOI:** 10.1101/2022.08.17.504168

**Authors:** Erithelgi Bertsoulaki, Hannah L. Glover, Joana I. Gomes-Neto, Barry Pizer, Helder Maiato, Sylvie Urbé, Michael J. Clague

## Abstract

We have identified USP31 as a microtubule and centrosome associated deubiquitylase, depletion of which leads to an increase in individual cell mass and defective proliferation. We have examined its dynamics and impact during mitosis. GFP-USP31 associates with the mitotic and central spindles, its levels are increased 2-3-fold in prometaphase compared to asynchronous cells and it is dynamically phosphorylated in a CDK1 dependent manner. We find that USP31 depleted cells display altered spindle morphology and chromosomal segregation errors, whilst stable expression of a catalytically inactive form of USP31 leads to polyploidy. At prometaphase, levels of multiple CPC components are destabilised, most prominently INCENP. Under anaphase conditions, depletion of USP31 impairs the translocation of both endogenous and ectopically expressed INCENP to the spindle mid-zone, whilst expression of catalytically inactive USP31 results in multiple ectopic cleavage furrows. In summary, our data indicate a multifaceted regulatory role for USP31 during mitosis, with a profound impact on chromosomal passenger complex abundance, dynamics and function.

---

The scope and range of protein ubiquitylation rivals that of phosphorylation (Kim et al., 2011; Xu et al., 2010). It is kept in check by a family of ~100 enzymes collectively known as the deubiquitylases (DUBs) (Clague et al., 2013). They may act to stabilise proteins that are turned over by either proteasomal and lysosomal degradation, but can also regulate ubiquitin signaling pathways. Individual DUBs may be linked with varying stringency to specific substrates by processing ubiquitin chains of different linkage type, confinement to a specific organelle or macromolecular structure, and by direct interaction with substrate protein (Clague et al., 2019). The ubiquitin specific protease (USP) family (~50 members), is the largest sub-family of DUBs that, with some notable exceptions, are relatively non-discriminatory between chain-linkage types (Faesen et al., 2011; Komander et al., 2009). Many members have specific substrate protein recognition motifs residing outside their conserved catalytic domain or as inserts within the catalytic domain (Clague et al., 2013; Ye et al., 2009).

Systematic studies of DUB localisation using GFP-tagged constructs revealed a number of DUBs that localise to clearly identifiable compartments such as ER (USP19), mitochondria (USP30) and plasma membrane (USP6) and a collection which can translocate to sites of DNA damage (Nishi et al., 2014; Sowa et al., 2009; Urbe et al., 2012). We have previously identified GFP-USP21 as a microtubule and centrosome associated DUB (Heride et al., 2016; Urbe et al., 2012). The USP family member CYLD is also known to bind microtubules through a CAP-Gly domain and to promote their assembly (Gao et al., 2008). Several DUBs have been associated with centrosomes (USP9X, USP21, USP33) or with the basal body of primary cilia (USP21, USP9X, CYLD) (Douanne et al., 2019; Han et al., 2019; Li et al., 2013; Li et al., 2017; Urbe et al., 2012). Here we identify, USP31 as a third microtubule-associated DUB that also decorates the centrosome. USP31 is one of the least explored USP family members, for which we now present the first substantive characterisation. In common with the majority of USP family members, USP31 processes ubiquitin chains of varying topologies (Takahashi et al., 2020).

Since the discovery of concerted protein degradation during the cell cycle, the central role for ubiquitylation has been clear. Although a small fraction of α/β tubulin itself is ubiquitylated, its function and significance is poorly understood. At the centrosome, γ-tubulin is ubiquitylated by BRCA1, which appears to limit centrosome duplication in mammary cells (Starita et al., 2004). Attention has been focused on the ubiquitylation of accessory proteins that govern the behaviour of microtubule networks. A key co-ordinator of cell cycle progression is the anaphase promoting complex (APC), a multi-subunit E3 ligase, which triggers the degradation of proteins that would otherwise restrict cell cycle progression. The timing of this degradation is crucial and those factors that are bound to microtubules are protected from degradation immediately after APC activation in distinction to their cytosolic counterparts (Song et al., 2014). Another critical co-ordinator is the Chromosomal Passenger Complex (CPC), comprised of one copy each of Aurora B kinase, Survivin, Borealin/DasraB and INCENP. This complex localises to centromeres during early mitosis and transfers to the spindle mid-zone and cleavage furrow during late mitosis (Carmena et al., 2012). Ubiquitylation of CPC components profoundly influences this choreography. Lys63-linked ubiquitylation of Survivin promotes CPC association with centromeres, and its release can be mediated by the deubiquitylase USP9X (Vong et al., 2005). Aurora B is ubiquitylated by two distinct CUL3 containing E3-ligase complexes at different stages of mitosis (Maerki et al., 2009; Sumara et al., 2007). In *Xenopus laevis*, the AAA-ATPase CDC48/p97, in complex with co-factors Ufd1 and NpI4, has been proposed to extract ubiquititylated Aurora B from chromosomes thereby facilitating chromosome condensation upon exit from mitosis (Ramadan et al., 2007). We now describe USP31 as a novel regulator of CPC dynamics and stability, under the control of the mitotic kinase CDK1.

## Results

### USP31 is a microtubule localizing DUB

In an earlier study we systematically described the localisation of 66 GFP-tagged DUB family members, but that collection did not include USP31 (Urbe et al., 2012). We now show that GFP-USP31 transiently expressed in mammalian cells localises to microtubules. Accordingly, co-localisation with α-tubulin is lost by cold induced microtubule depolymerisation (**Fig. 1A**). GFP-USP31 can also be found to co-localise with Pericentrin at the centrosome and with ARL13b at primary cilia (**Fig. S1A**). We set about mapping the region of USP31 responsible for microtubule association. USP31 contains a catalytic domain, which, at the primary sequence level, is subject to an extensive insertion that includes a ubiquitin-like (Ubl) domain (aa 314-402; **Fig. 1B**). C-terminal to the catalytic domain lies a long Serine rich (Ser-rich) region that starts with a predicted SxIP motif (aa 765-774). Such a motif, set within Ser-rich sequence stretches, is commonly found in microtubule +end tracking proteins (+TIPs) where it mediates the association with end binding proteins, like EB1 and EB3 (Honnappa et al., 2009). Here we show that the Ser-rich C-terminal region is necessary and sufficient for microtubule localisation but the SxIP motif appears to be dispensable (**Fig. 1B** and **Fig. S1B,C**).

**Fig. 1:**
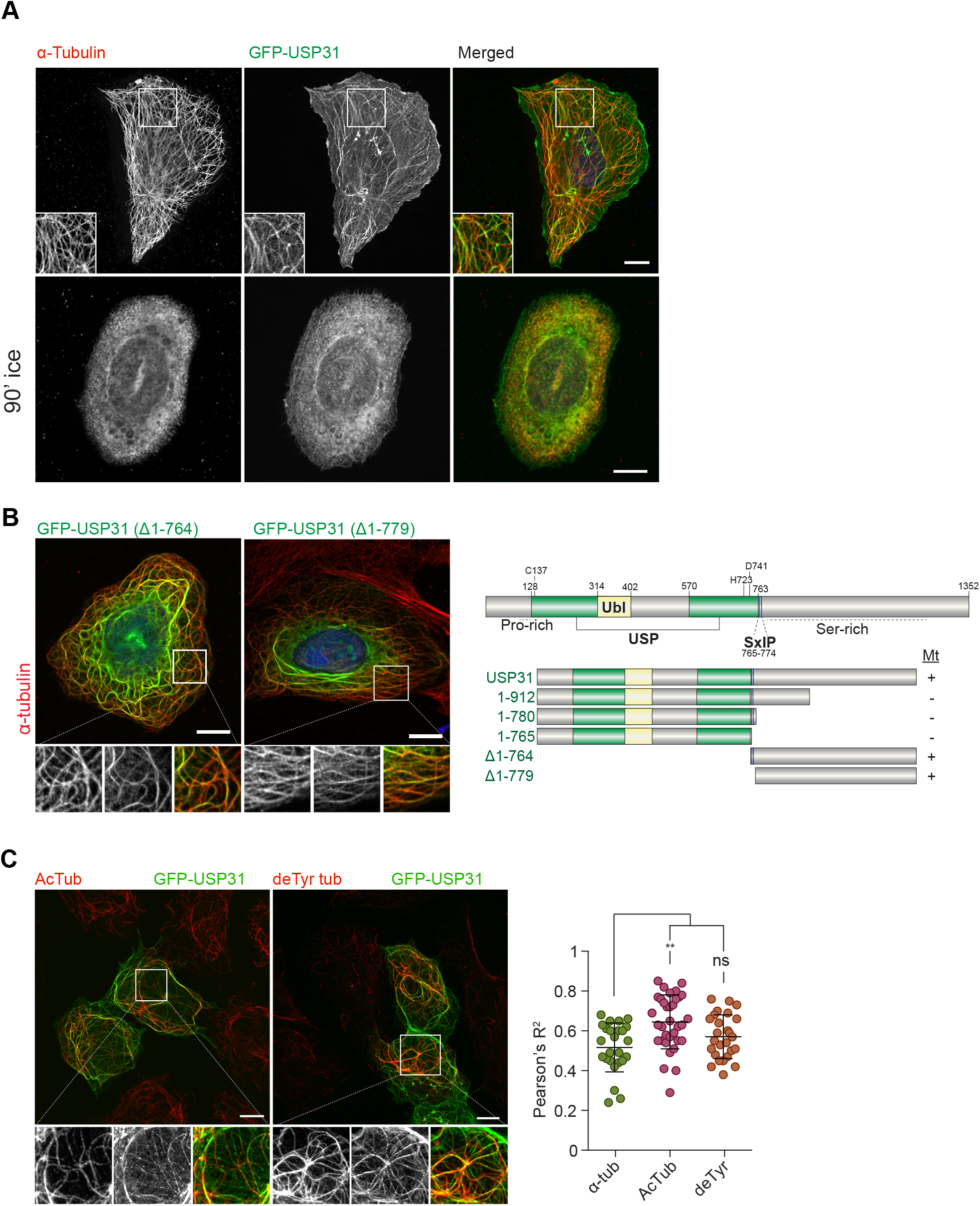
USP31 is a microtubule localising DUB. **A.** U2OS cells were transiently transfected with GFP-USP31. Microtubules were cold-depolymerised (90’ min on ice) or not, prior to MeOH fixation, and cells stained for α-tubulin. Images were acquired on a 3i spinning disk confocal microscope. Scale bar: 10 μm. **B.** The C-terminal region of USP31 is necessary and sufficient for microtubule association. Representative images and schematic representation of the truncated constructs used to map the microtubule localising region. Scale bar: 10 μm. Ubl: Ubiquitin-like domain. **C.** USP31 localises to post-translationally modified microtubules. U2OS cells were transfected with GFP-USP31, fixed with MeOH and stained for acetylated (AcTub) and detyrosinated (deTyr tub) tubulin. Pearson’s correlation coefficient was calculated using the Coloc2 plugin of Fiji. Shown is the average ±SD for at least 25 cells (α-tub: 25, AcTub: 34, deTyr: 26). Kruskal-Wallis test was used with Dunn’s multiple comparison test; **p≤0.01, ns: not significant.

We noticed that only a subset of microtubule structures is decorated with GFP-USP31 and sought to examine co-localisation with different forms of post-translationally modified tubulin. We observed an increased overlap between acetylated (K40) tubulin and GFP-USP31 compared to total or detyrosinated α-tubulin populations (**Fig. 1C**). In interphase cells, this suggests a link to a longer lived fraction of microtubules (Janke and Montagnac, 2017).

### USP31 depletion and inactivation causes mitotic defects

Western blot analysis of a cell line panel shows that USP31 expression is widespread (**Fig. S1D**). We focused our analysis on U2OS cells due to their high suitability for imaging based experiments. We were unable to obtain viable CRISPR KO clones but we could efficiently deplete USP31 with multiple siRNA oligonucleotides (Q1, Q2, Q4; **Fig. S1E**). Consistently we found larger U2OS cells following USP31 depletion as judged by an increase in the total cell and nuclear surface area (**Fig. 2A**). Critically, this was not simply a manifestation of enhanced cell spreading, but was reflected in increased cellular protein mass (**Fig. 2A**). No gross disruption of the α-tubulin stained interphase microtubule network was evident. Flow cytometry revealed a significant decrease in G1 and a clear increase in G2/M populations for two out of three siRNAs targeting USP31, without a clear indication for growth arrest at any particular phase of the cell cycle (**Fig. 2B**). By continuously imaging U2OS cells stably expressing mRFP-H2B, we observed that siRNA depletion of USP31 led to a reduction in the number of cells entering mitosis (**Fig. 2C**, left panel). For those cells that did enter mitosis, we assessed the time taken to progress from the point of chromatin condensation to onset of anaphase. It is evident that loss of USP31 increases the time individual cells spend in mitosis (**Fig. 2C**, right panel).

**Fig. 2:**
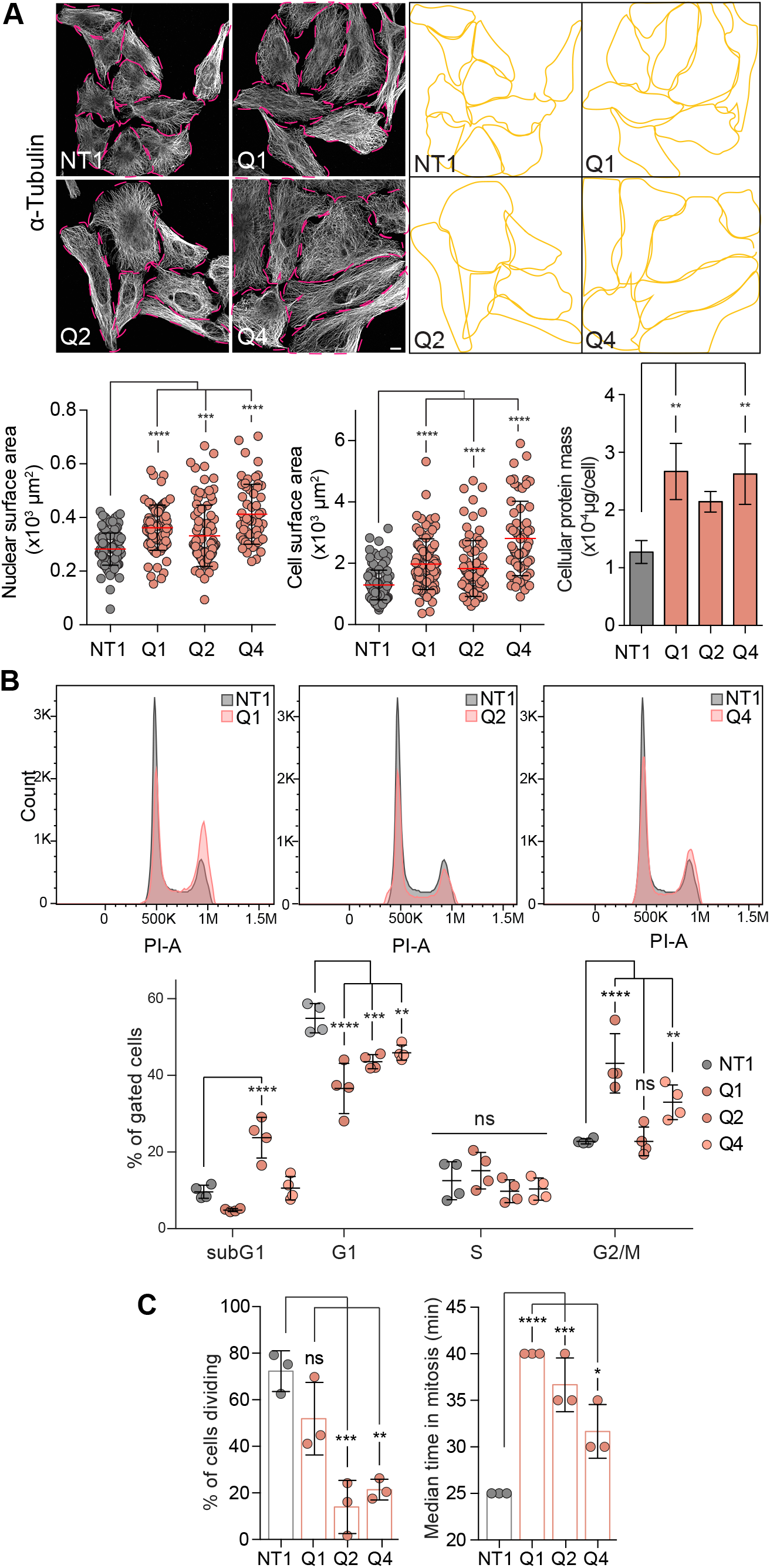
Depletion of USP31 causes mitotic defects. **A.** USP31 depletion causes an increase in cell and nuclear surface area. U2OS cells were transfected with either control (NT1) or three siRNAs targeting USP31 (Q1, Q2, Q4) for 72 h, fixed with MeOH and stained for α-tubulin. The cell and nuclear surface area were determined by drawing the perimeter around each cell and nucleus using ImageJ. Shown are the individual data from at least 57 cells per condition (n=2 independent experiments) as well as the average ±SD. Right hand bar shows cellular protein mass from three independent experiments; average ±SD. (Cell count for cell surface area: NT1:127, Q1:66, Q2:89, Q4:57; cell count for nuclear surface area: NT1:139, Q1:89, Q2:81, Q4:61). Oneway ANOVA test with Dunnet’s multiple comparison was used. **B.** Cell cycle profile of U2OS cells depleted of USP31 for 72 h. Cells were fixed and stained with propidium iodine before subjected to flow cytometry. Representative graph of the cell cycle profile of the gated cells are shown. Bottom Graph shows the distribution of the cells in 4 independent experiments (error bars show SD; two-way ANOVA with Bonferroni’s multiplex comparisons test. **C.** USP31 depletion delays mitotic progression. mRFP-H2B U2OS cells were depleted of USP31 for 48 h. They were then imaged over 16 h and monitored throughout mitosis using a Nikon Ti-Eclipse microscope. Shown are the percentage of cells that divided during this time as well as the time cells spent from nuclear envelope breakdown to anaphase (n=3 independent experiments; error bars show SD; one-way ANOVA with Bonferroni’s multiplex comparisons test. **p≤0.01, ***p≤0.001, ****p≤0.0001

We generated U2OS cells stably expressing GFP-USP31 (WT) or a catalytically inactive mutant (C137A; CA), which can in principle act in a dominant negative fashion. Both of these proteins localise to microtubules and centrosomes (**Fig. S2A, B**). No consistent changes in growth between multiple clonal lines were seen (**Fig. 3A**). This most likely reflects that compensation of any major growth defect is a requirement for isolation of the clones. However, cells expressing inactive GFP-USP31(C137A), but not wild-type GFP-USP31, are substantially larger, recapitulating the increased cell area and cell mass associated with USP31-depleted cells (**Fig. 3B**). Interestingly, FACS analysis reveals a tetraploid status for cells expressing catalytically inactive USP31 (CA1), indicative of impaired cytokinesis or mitotic slippage (**Fig. 3C**).

**Fig. 3:**
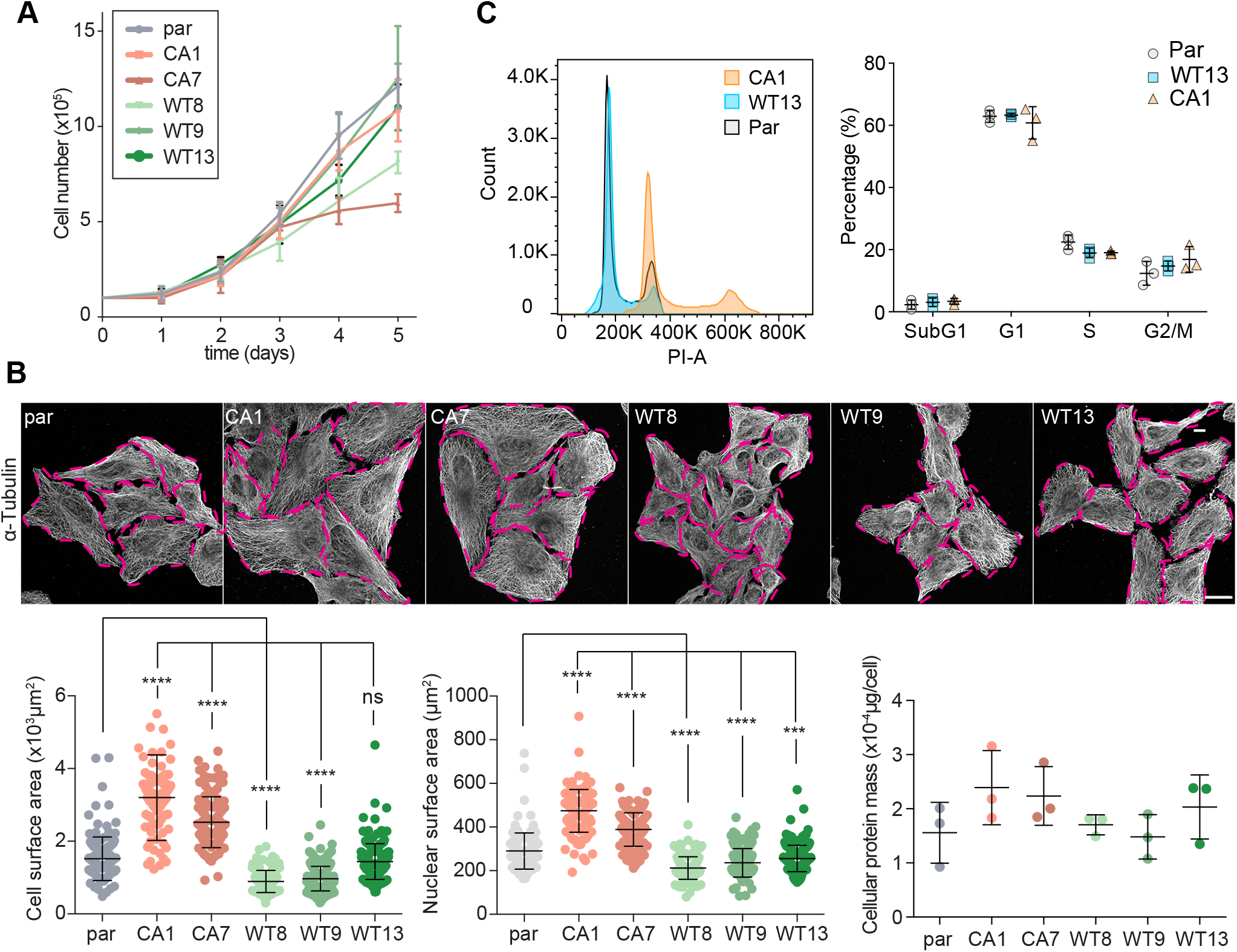
Mitotic phenotype analysis of stable cell lines overexpressing either wild-type or catalytically inactive USP31. **A.** Growth curve of parental U2OS cells (par) or cells stably expressing wild-type (WT8, WT9, WT13) or catalytically inactive (CA1, CA7) USP31 (n=3 independent experiments, error bars show SD). **B**. Stable overexpression of catalytically inactive USP31 induces an increase in cell size and protein mass. Parental U2OS cells (par) or cells stably expressing wild-type (WT8, WT9, WT13) or catalytically inactive (CA1, CA7) USP31 were fixed with MeOH and stained for α-tubulin. The cell and nuclear surface area was determined by drawing around each cell using ImageJ (parental: 137, CA1: 102, CA7: 120, WT8: 152, WT9: 234, WT13: 184) and nucleus (parental: 115, CA1: 121, CA7: 130, WT8: 164, WT9: 163, WT13: 164). Shown are data from >100 cells per condition and from two independent experiments, error bars show SD; one-way ANOVA with Dunnett’s multiple comparison test). ***p≤0.001, ****p≤0.0001. Cellular protein mass was also calculated for the parental and stable cell lines (n=3 independent experiments; error bars show SD). **C.** Cell cycle profile of the parental U2OS cells and the stable cell lines. Cells were fixed and stained with propidium iodine before subjected to flow cytometry. Representative graph of the cell cycle profile of the gated cells are shown (Par - 48160; WT13 - 45915; CA1 - 48237). Graph shows the distribution of the cells in 3 independent biological experiments; error bars show SD.

### USP31 localises to the spindle and its depletion increases kinetochore microtubule turnover

Due to the cell cycle defects observed upon USP31 depletion, we decided to further investigate its behaviour and role in mitosis. Imaging GFP-USP31 dynamics during mitosis showed only a minor fraction associated with the metaphase spindle. However, upon anaphase onset, we observed a rapid recruitment onto kinetochore microtubules and onto the central spindle (**Fig. 4A**, **Video 1**). Upon USP31 depletion, defects in spindle organisation were apparent, with metaphase spindles becoming more elongated and asymmetric (**Fig. 4B**). We next enquired if USP31 can influence microtubule dynamics during mitosis, utilising a U2OS cell line that stably expresses both Cherry-tubulin and photoactivatable PA-GFP-tubulin. We arrested cells in metaphase using a proteasome inhibitor (MG132) and generated a stripe of fluorescent tubulin in one half-spindle next to the metaphase plate. Z-stacks were acquired at 15-second intervals and the fluorescence dissipation was analysed as previously described (Girao and Maiato, 2020). Depletion of USP31 led to a clear increase in kinetochore microtubule (kMT) turnover indicated by a decrease in kMT half-life (**Fig. 4C** and **D**). The half-life of non-kinetochore microtubules was not affected nor was the ratio of kinetochore to non-kinetochore microtubules (**Fig. 4D**). Consequently, the high turnover of kinetochore microtubules was associated with chromosomal segregation errors in anaphase. These include a significant increase in cells with multiple lagging chromosomes in anaphase (**Fig. 4E**).

**Fig. 4:**
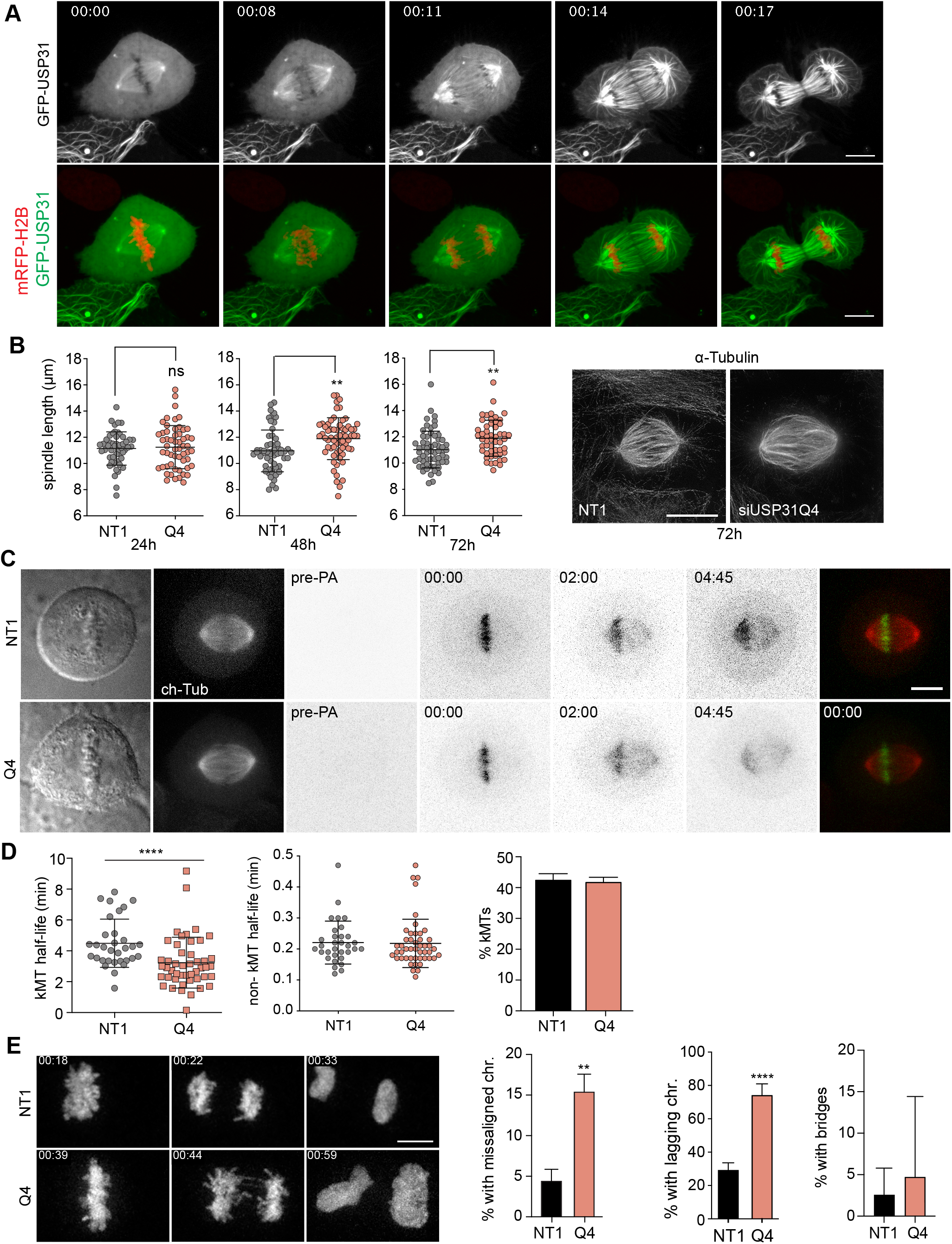
USP31 localises to the spindle and its depletion causes spindle distortion and alters kinetochore microtubule dynamics. **A.** U2OS cells stably expressing mRFP-H2B were transfected with GFP-USP31 and a z-stack was acquired at specified time points using a 3i spinning disk confocal microscope. The maximum projection is shown. Scale bar = 10 μm. Time is shown in hours:min. See also Video 1. **B.** U2OS cells were transfected with either control (NT1) or an siRNA targeting USP31 (Q4) for either 24 h, 48 h or 72 h, then fixed with MeOH, stained for α-tubulin and DAPI and imaged on a Mariana 3i spinning disk confocal (for measurements) or a using an Abberior ‘Expert Line’ gated-STED microscope (image shown). Scale bar = 10 μm. Metaphase cells (> 50 per condition) were selected, and their spindle length calculated. Cell numbers for 24 h, 48 h and 72 h respectively for NT1: 51, 56, 60 and for Q4: 56, 64, 52). One-way ANOVA with Bonferroni’s multiplex comparisons test, **p≤0.01. **C.** U2OS cells stably expressing PA-GFP-tubulin and Cherry-tubulin were transfected with siRNA targeting USP31 (Q4) or Non-targeting siRNA (NT1) for 24 h. Cells were treated with MG132 (5 μM) for 1 h prior to imaging. Metaphase cells were selected based on the DIC image, and a pre-photoactivation image was acquired. In the panel are shown: the DIC image, the Cherry-tubulin (ch-Tub) image, the PA-GFP-tubulin signal before (pre-PA) and the first frame after photoactivation (00:00), as well as time-points of 2.00 and 4.45 min after photoactivation. Scale bar = 5 μm. **D.** The kinetochore microtubule half-life for cells treated as in C was calculated for 32 (NT1) and 47 (Q4) cells. (Data shown are from 8 independent experiments; Mann-Whitney U test ****p≤0.0001). The non-kinetochore microtubule half-life percentage of kinetochore microtubules are also shown, error bars indicate SD. **E.** Chromosome congression and segregation fidelity is compromised upon USP31 depletion. U2OS cells stably expressing GFP-H2B and Cherry-tubulin were transfected with control (NT1) or USP31 targeting siRNA (Q4) for 48 h, then imaged on a modified Nikon TE2000-E based spinning disk confocal microscope and assessed for chromosome fidelity errors. Shown is a maximum projection. Scale bar = 5 μm. The percentage of cells entering anaphase with misaligned chromosomes (chr.) as well as cells with lagging chromosomes and bridges are indicated in the graphs (NT1: 87 cells, Q4: 86 cells; from n=4 independent experiments, unpaired t-test for misaligned chromosomes, Two-way ANOVA with Šídák’s multiple comparisons test for others; **p≤0.01, ****p≤0.0001).

### USP31 regulation during mitosis

We next examined protein expression levels of USP31 throughout the cell cycle using standard synchronisation procedures (**Fig. 5A**). Total USP31 levels increase on average two to three-fold as cells enter mitosis and remain elevated throughout (**Fig. 5B,C** and **Fig. S3A** and **B**). We also noticed that this increase was accompanied by a slower migration in SDS-PAGE gels, which we surmised could represent phosphorylation. To assess this, we treated cell lysates arrested in prometaphase with lambda phosphatase, which restored the higher electrophoretic mobility of the ~150kD full length form and a lower MW form running at ~100kD, which may correspond to a splice variant (**Fig. 5D**, blue and green arrowheads respectively). This confirms that USP31 is indeed phosphorylated during mitosis. Furthermore, the kinetics of its accumulation/phosphorylation upon entry into mitosis (**Fig. S3A**) and depletion/dephosphorylation as the cells progress through mitosis (**Fig. S3B**) closely mirror those of CyclinB and CDC27 respectively.

**Fig. 5:**
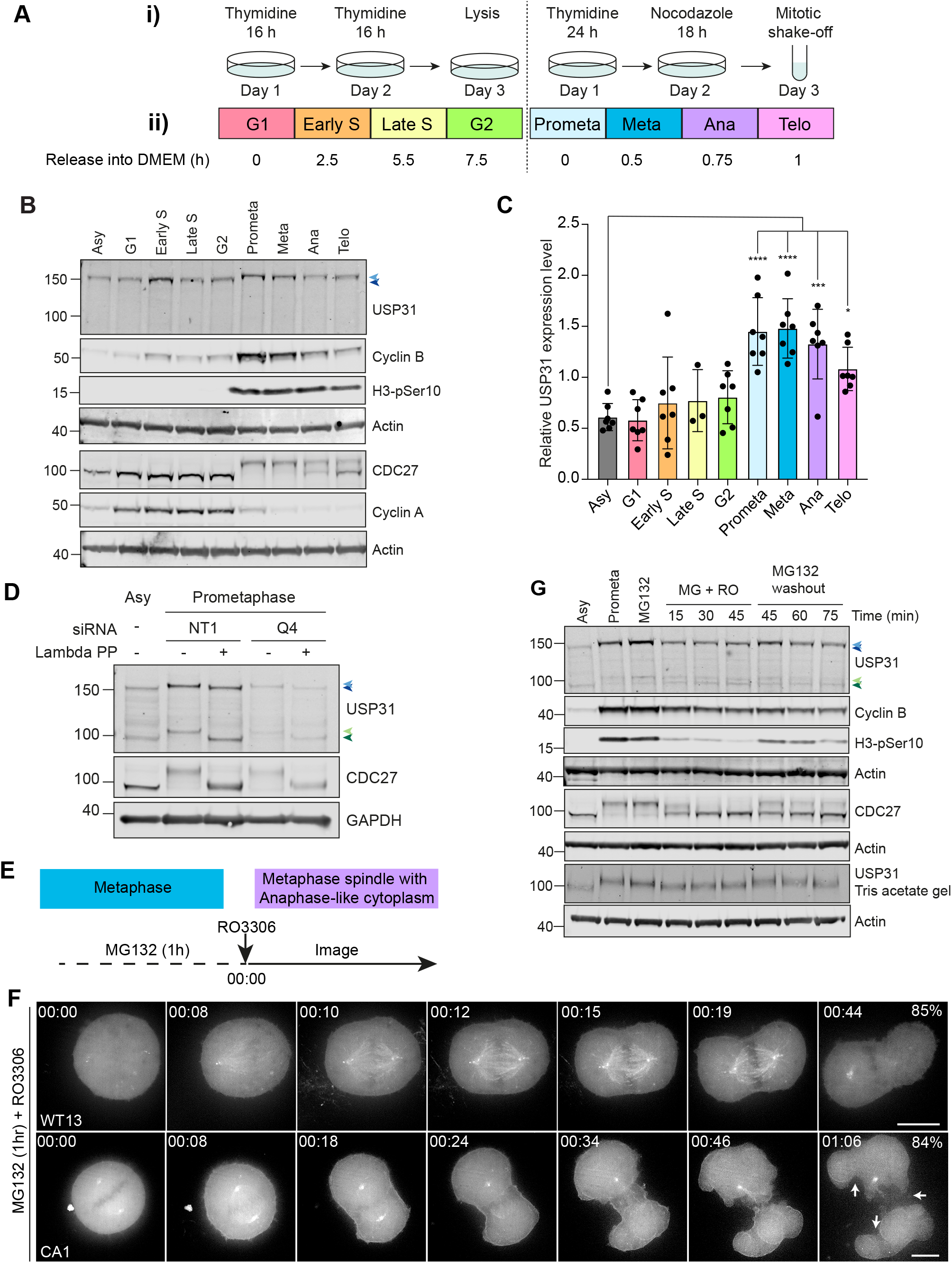
USP31 regulation during mitosis. **A.** Schematic showing cell synchronisation protocols, separated by a dashed line. **i)** schedules for double thymidine block or single thymidine block followed by nocodazole treatment are shown. **ii)** Lysis schedule for each stage is shown. **B.** USP31 expression during the cell cycle. U2OS urea cell lysates were collected upon either synchronisation protocol at the time-points specified in A. Dark blue arrow shows the unphosphorylated band, light blue arrow shows the phosphorylated. **C:** Western blot quantitation of **B**. Graph shows individual values from 3 (Late S) or 7 (all others) independent biological repeats. Error bars indicate SD; one-way ANOVA with Dunnett’s multiple comparisons test, *p≤0.05, ***p≤0.001, ****p≤0.0001 **D.** U2OS cells were transfected with either control (NT1) or an siRNA oligos targeting USP31 (Q4) for 48h and synchronised using a single thymidine block followed by nocodazole treatment. Cells were lysed and samples were treated with 400 units of lambda phosphatase for 30 min. Blue and green arrows indicate full length USP31 and a proposed shorter isoform, respectively. Top arrows (light) and bottom arrows (dark) indicate the phosphorylated and unphosporylated USP31 species respectively. **E.** Schematic showing the experimental protocol for cells shown in F. **F.** Localisation of USP31 at metaphase is controlled by CDK1-dependent phosphorylation. U2OS cells stably expressing wild-type (WT13) or catalytically inactive (CA1) USP31 were arrested in metaphase using MG132 (5 μM) for 1 h. Cells were imaged immediately after the addition of RO3306 (CDK1 inhibitor; 10 μM) using a 3i-spinning disk confocal. Z-stacks were acquired every minute, and the maximum projection for the selected time-points are shown (hours:mm). Arrows point to furrow positions in the CA1 cell line. Scale bar: 10 μm. Time is shown in hours:min. See also Video 2 and 3. **G.** U2OS cells were synchronised via single thymidine block followed by nocodazole and released into fresh DMEM. One sample was lysed immediately (Prometa), whereas MG132 (MG, 5 μM) was added to all other samples for 1 h to arrest cells at the metaphase. CDK1 inhibitor (RO3306, RO, 10 μM) was then added to the other samples and incubated for specified times. For control cells, MG132 washout in fresh DMEM was performed and cells incubated for specified times to allow re-entry into mitosis. Asy: Asynchronous cell lysates. Blue and green arrows indicate full length USP31 and a proposed shorter isoform, respectively. Top arrows (light) and bottom arrows (dark) indicate the phosphorylated and unphosporylated USP31 species respectively.

We noticed that the rapid recruitment of USP31 to spindle microtubules during anaphase (**Fig. 4A, Video 1**), coincides with its apparent dephosphorylation and speculated that it is the loss of CDK1 activity that initiates this relocalisation. We next blocked cells in metaphase using the proteasome inhibitor MG132, and then created an anaphase-like cytosol by inhibiting CDK1 with RO3306, all the while maintaining a metaphase spindle (**Fig. 5E**). We observed a rapid accumulation of both wild-type and catalytically inactive GFP-USP31 at the arrested metaphase spindle (**Fig. 5F**). Biochemical analysis of lysates corresponding to these conditions indicated the dependence of USP31 phosphorylation on CDK1 activity. Treatment with RO3306 leads to rapid dephosphorylation similar to MG132 wash-out and allows cells to progress through mitosis, albeit with slower kinetics (**Fig. 5G** and **Fig. S3C, D, E**). Interestingly, inhibition of Aurora B at the same point (with ZM447439) did not change the phosphorylation status of USP31 (**Fig. S3C, D, F**). Thus, we conclude that the reduction of CDK1 activity is responsible for both the dephosphorylation of USP31 and its recruitment to the kinetochore microtubules and the central spindle upon anaphase onset.

### USP31 modulates CPC abundance and localisation during mitosis

In this anaphase-like condition, we observed a peculiar phenotype in the cell lines that were stably expressing catalytically inactive GFP-USP31(C137A). In these cells, inhibiting CDK1 leads to ectopic furrowing of the cytoplasm at multiple locations (**Fig. 5F**; 38/45 – 84% cells for the CA mutant compared to 6/40 – 15% for the WT cells; **Video 2 and 3**). We were intrigued by this observation, and we speculated a possible involvement of the CPC complex, as this is the major governor of furrow positioning (Carmena et al., 2012). Both parental and cells over-expressing the wild-type enzyme showed accumulation of the CPC component Aurora B at a single furrow site location as expected. In cells overexpressing catalytically inactive USP31(C137A), Aurora B was localised to each of the multiple furrowing positions (**Fig. 6A**).

**Fig. 6:**
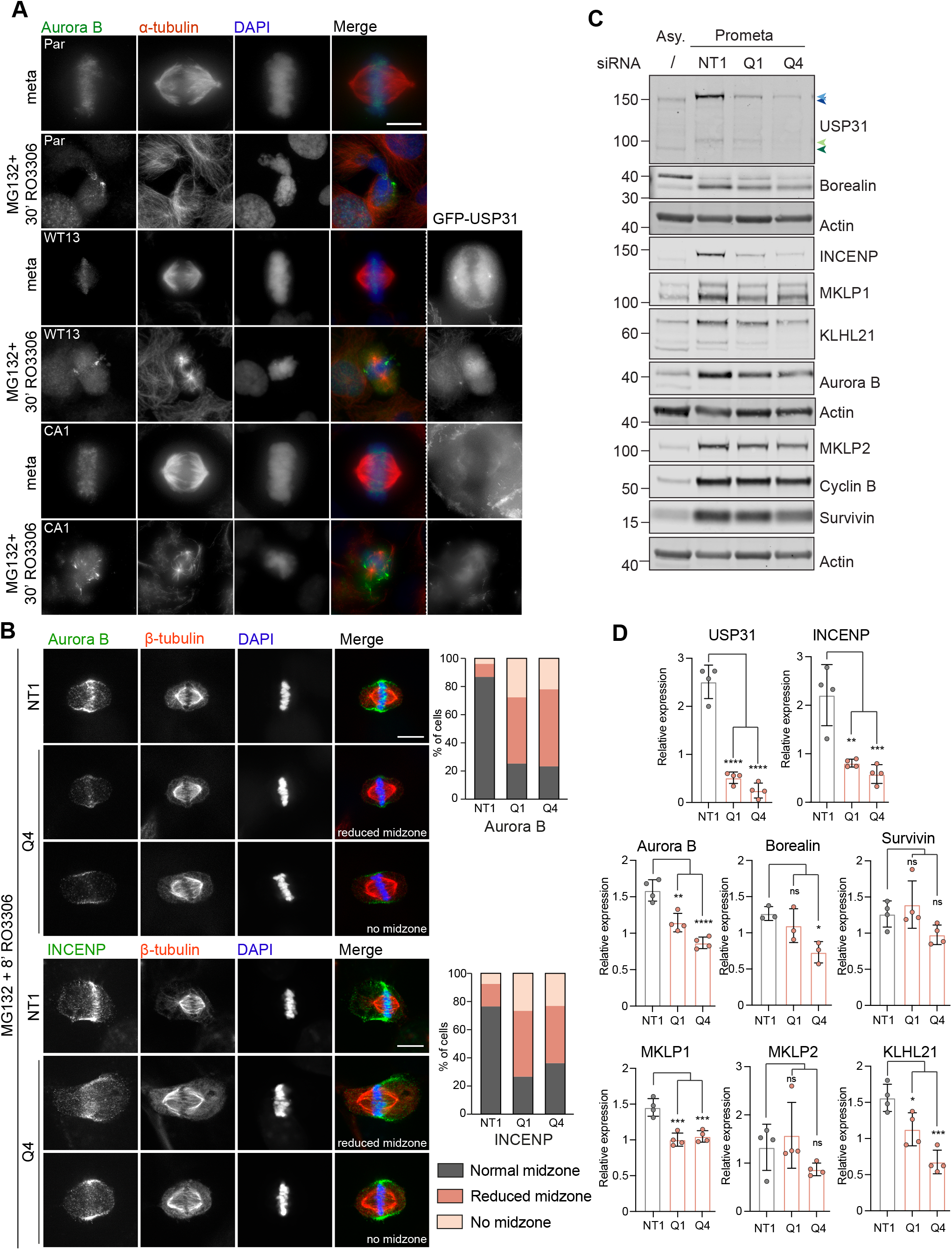
USP31 modulates CPC stabilisation and localisation during mitosis. **A.** Parental U2OS cells (Par), as well as the stable cell lines overexpressing USP31 wild type (WT13) and catalytically inactive (CA1), were arrested in metaphase with MG132 (5 μM) for 1 h (meta) and either fixed immediately or further treated for 30 min with RO3306 (10 μM) prior to fixation with MeOH and staining for the indicated proteins. Images were acquired with a Zeiss AxioImager Z1 microscope. Scale bar = 10 μm. **B.** U2OS cells were transfected with siRNA against USP31 (Q4) or a non-targeting control (NT1) for 48 h, then treated with MG132 (5 μM) for 1 h and subsequently treated with RO3306 (10μ M) for 8 min. Cells were fixed with MeOH and stained for the indicated antibodies. Images were acquired with a 3i spinning disk confocal; a single confocal slice is shown. Scale bar 10 μM. Bar graph show the quantification of the frequency of phenotypes observed (% cells showing each phenotype; cells analysed per condition: Aurora B: NT1 (76), Q1 (83), Q4 (73). INCENP: NT1 (119), Q1 (109), Q4 (160)). **C.** U2OS cells were transfected with siRNA against USP31 (Q1 and Q4) or a non-targeting control (NT1) for 48 h. Cells were synchronized to prometaphase (Prometa) using thymidine and nocodazole and then lysed immediately. Samples were analysed by western blotting and probed for indicated proteins. Asy: Asynchronous cell lysates. Shown is a representative experiment. **D.** Quantification of western blots illustrated in C. Graph shows results from 4 independent experiments. Statistical analysis carried out via one-way ANOVA with Dunnett’s multiple comparisons test. ns=not significant, *p≤0.05, **p≤0.01, ***p≤0.001, ****p≤0.0001.

At early anaphase, the CPC associates with the mitotic Kinesin-like protein MKLP2 and undergoes a dramatic relocalisation from the centromeres to the spindle mid-zone and to the equatorial cortex (cell periphery), defining the position of the cleavage furrow (Adriaans et al., 2020; Gruneberg et al., 2004). In cells depleted of USP31, multiple components of the CPC complex (INCENP, and Aurora B) fail to efficiently translocate to the mid-zone, whilst the cortical (peripheral) localisation is less affected (**Fig. 6B** and **Fig. S4A** and **B**). We next examined the expression levels of CPC proteins and associated carriers upon USP31 depletion in mitotic cells. Many of these proteins, in particular INCENP, Aurora B as well as the centralspindlin component, MKLP1, show clear reductions in their levels (**Fig. 6C** and **D**). A corresponding reduction in INCENP but not Aurora B can also be seen in cells overexpressing catalytically inactive USP31(C137A) but not wild-type USP31 (**Fig. S4C** and **D**). In principle, such a decrease in protein expression levels could be due to either reduced transcription or enhanced protein turnover. However, we can recapitulate the impact of USP31 depletion on INCENP expression levels in doxycycline inducible INCENP-GFP expressing U2OS cells, arguing against a transcriptional mechanism and supporting a direct role of USP31 in stabilising INCENP (**Fig. 7A**). These cells also allowed us to monitor the dynamics of ectopically expressed INCENP-GFP in asynchronous cells. In addition to a marked decrease in intensity, we observed a clear delay in the translocation from the centromeres to the midzone (**Fig. 7B**, **Video 4-6**). We conclude that USP31 influences both abundance and dynamics of the CPC.

**Figure 7:**
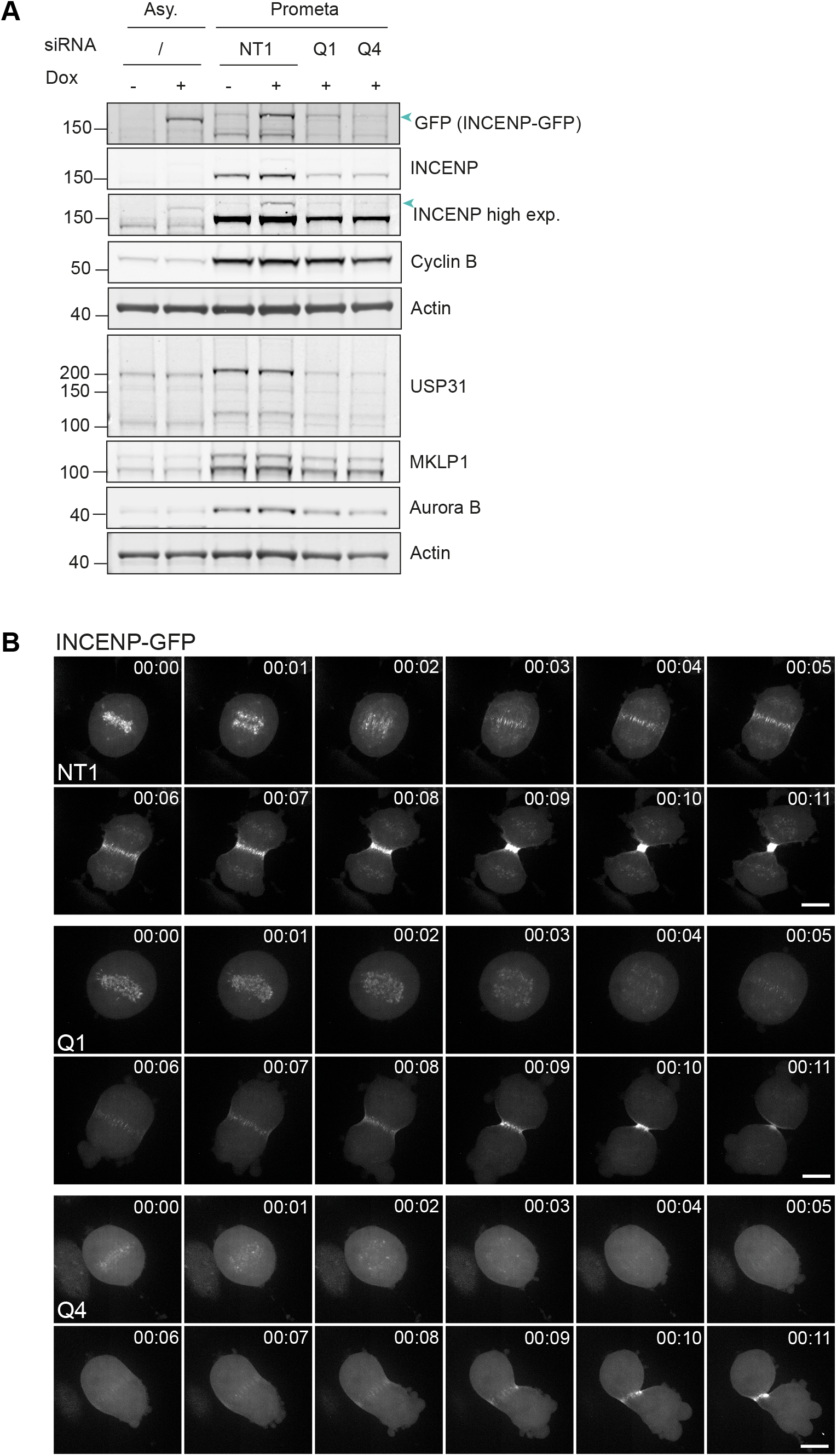
USP31 depletion destabilises ectopically expressed INCENP and delays its translocation to the midzone. **A.** U2OS cells inducibly expressing INCENP-GFP were transfected with siRNA against USP31 (Q1, Q4) or a non-targeting control (NT1) and synchronised with a single thymidine block followed by nocodazole to prometaphase. INCENP-GFP (arrowheads) expression was induced for 3 h with doxycycline (1 μg/ml) prior to lysis and processing for western blotting with the indicated antibodies. **B.** U2OS cells inducibly expressing INCENP-GFP were transfected with siRNA against USP31 (Q1, Q4) or a non-targeting control (NT1) for 45 h, then treated with doxycyclin (1 μg/ml) for 3 h to induce expression of INCENP-GFP. Cells were then imaged on a 3i spinning disc confocal microscope as they transition from metaphase to anaphase. Z-stacks (range 17μm, 1μm steps) were acquired every minute. Shown is a sequence of representative Z-stack maximum projections starting one frame before anaphase onset. Scale bar = 10 μm. Time is shown in hours:min. See also Video 4,5,6.

## Discussion

We have provided a foundational study of the hitherto poorly characterised DUB USP31, which joins a small collection of DUBs that localise to microtubules. Whilst we were unable to detect endogenous USP31 levels by immunofluorescence microscopy with currently available antibodies, our data using GFP-tagged protein suggest a preferential association with acetylated microtubules. Its cell cycle-dependent expression levels, phosphorylation status and localisation dynamics all indicate a role during cell division. Accordingly, we have shown that multiple features of mitotic progression fall under the influence of USP31 activity.

The destabilisation of kinetochore microtubules and associated segregation errors, as well as elongated and distorted spindle morphology in USP31 depleted cells, indicate an impact on microtubule dynamics that is yet to be fully explored elsewhere. Likewise, it is currently unclear whether CDK1 dependent phosphorylation controls USP31 activity and/or microtubule association. Importantly, at least in part, the observed delay in mitotic progression upon USP31 depletion can be attributed to the regulation of the CPC.

We observed a clear decrease in multiple CPC components at prometaphase in both USP31-depleted and catalytically inactive mutant USP31 expressing cell lines that was most profound for INCENP. We show that ectopically expressed INCENP is equally sensitive to USP31 depletion, supporting a post-translational regulatory mechanism. Combining all of our findings, the most parsimonious explanation is that USP31 acts locally to suppress ubiquitylation of CPC components, particularly INCENP, impacting upon its stability. In addition, the extraction of CPC components from chromosomes is known to be under the control of a cycle of ubiquitylation and deubiquitylation (Maerki et al., 2009; Ramadan et al., 2007; Sumara et al., 2007; Vong et al., 2005). Our finding that this migration is delayed upon USP31 depletion suggests a second role in this process.

Recently, DUBs have emerged as a druggable class of protein targets and several highly specific inhibitors of individual USPs have been introduced (Clancy et al., 2021; Gavory et al., 2018; Kategaya et al., 2017; Lamberto et al., 2017; Rusilowicz-Jones et al., 2022; Turnbull et al., 2017). We believe that the work presented here will provide a springboard for further investigation of USP31 and pursuit of its interesting biology with the prospect of translational applications.

## Materials and Methods

### Cell lines

U2OS from the European Collection of Authenticated Cell Cultures (ECACC), as well as U2OS derived cell lines, and NIH3T3 cells were grown in Dulbecco’s Modified Eagle Medium (DMEM), and hTERT-RPE1 cells in DMEM/F12, each supplemented with 10% heat-inactivated foetal bovine serum in 5% CO2. U2OS cells stably overexpressing mRFP-H2B, and U2OS cells expressing PA-GFP-α-tubulin/mCherry-tubulin were described in (Barisic et al., 2014; Ferreira et al., 2018; Ferreira et al., 2020). U2OS cells with doxycycline inducible VSV-INCENP-GFP expression were a kind gift of Dr. Susanne Lens (Utrecht University; (van der Waal et al., 2012)). To generate USP31 stably overexpressing cell lines, U2OS cells were transfected with GFP-USP31WT(Q2R) and GFP-USP31CA(Q2R) (C137A; catalytically inactive mutant) using Genejuice, grown under selection with G418 (0.4 mg/ml) for two weeks, single cell diluted, screened for GFP-USP31 expression and amplified.

### Antibodies

Primary antibodies used: Actin mouse (Proteintech, 66009-1-Ig), Actin rabbit (Proteintech, 20536-1-AP), Aurora B (BD Transduction, 611083), Borealin (Santa Cruz, sc-376635), Cdc27 (BD Transduction, 610454), Cyclin A2 (Abcam, ab32386), Cyclin B1 (Abcam, 05-373), GAPDH (Cell Signaling, 2118S), GFP (Made in house), Histone H3 Phospho-Ser10 (Millipore, 05-806), INCENP (Santa Cruz, sc-376514), KLHL21 (Proteintech, 16952-1-AP), MKLP1 (Abcam, ab174304), MKLP2 (Bethyl, A300-879A), Survivin (Novus Biologicals, NB500-201), mouse α-tubulin (Sigma, T6199), rat α-tubulin (Biorad, MCA77G), β-tubulin (Abcam, ab6046), mouse acetylated α-tubulin (Sigma, T6793), rabbit detyrosinated α-tubulin (Abcam, ab48389), mouse γ-tubulin (Sigma (T5326), rabbit Pericentrin (Abcam, ab4448), USP31 (Santa Cruz, sc-100634). Secondary antibodies used for immunoblotting obtained from LICOR Biosciences: Donkey anti-mouse IRDYE 800CW (926-32212), Donkey anti-mouse IRDYE 680CW (926-32222), Donkey anti-rabbit IRDYE 800CW (926-32213), Donkey anti-rabbit IRDYE 680CW (926-32223), Donkey anti-sheep IRDYE 800CW (926-32214), Donkey anti-sheep IRDYE 680CW (926-32224). Secondary antibodies used for immunofluorescence obtained from Invitrogen: Donkey anti-mouse AF350 (A10035), Donkey anti-rabbit AF350 (A10039), Donkey anti-rabbit AF488 (A21206), Donkey anti-rabbit AF594 (A21207), Donkey anti-mouse AF488 (A21202), Donkey anti-mouse AF594 (A21203), Donkey anti-rat AF594 (A21209).

### Plasmids

Full length USP31 (aa 1-1352) and USP31 truncations (aa 1-765, aa 1-780, aa 1-912, aa Δ1-764 and aa Δ1-779) were amplified by PCR from pENTR223.1-USP31_BC156334 (Source Bioscience, IMAGE ID 100061896) and subcloned into the BglII and SalI sites of the pEGFP-C1 vector. The C137A (CA) mutation was introduced in USP31 by site-directed mutagenesis and the following primers: (5’-caaccacggcaacacgGCAttcatgaacgccacgc-3’ and 5’-gcgtggcgttcatgaaTGCcgtgttgccgtggttg-3’) and subsequently subcloned into pEGFP-C1 as described above.

### Cell lysis, SDS-PAGE and Western blots

Cells were placed on ice, rinsed 2 times with ice-cold PBS and lysed in urea lysis buffer (8M Urea, 50mM Tris-HCl pH8) unless specified. Samples were briefly sonicated before insoluble material was removed by centrifugation at 13,500rpm for 15 min at 4°C, and the protein concentration of the supernatant assessed by Pierce 660nm protein assay reagent (Sigma, 22660). All samples were diluted to a final 1X Laemmli sample buffer (40% glycerol, 8% SDS, bromophenol blue, 200 mM Tris base pH 6.8), boiled (95°C, 5 min) and proteins separated alongside protein standards (Rainbow and Broad Range) at 180V on precast either NuPAGE Novex 4-12% Bis-Tris Gels in 3-(N-morpholino)propanesulfonic acid (MOPS) buffer (Invitrogen), or NuPAGE™ 3-8% Tris-Acetate gels in NuPAGE™ Tris-Acetate SDS Running Buffer (Invitrogen) where specified. Samples were transferred to 0.45 μm nitrocellulose membrane (Amersham™ Protran^®^) (0.9A, 1-1.5 h) in transfer buffer (0.2 M Glycine, 25 mM Tris, 20% methanol). Membranes were stained with Ponceau S to confirm successful transfer, then incubated for 1 h in blocking solution (Tris buffered saline [TBS: 100 mM NaCL, 10 mM Tris, pH 7.5] supplemented with 0.1% Tween-20 [TBST, v/v], and 5% Marvel [w/v]), prior to incubation with specified primary antibodies overnight in a coldroom. Membranes were washed 3X for 5 min in TBST before incubation with IRDye-conjugated anti-mouse, anti-rabbit or anti-sheep secondary antibodies in blocking solution for 1 h and visualised using an Odyssey infrared scanner (LI-COR Biosciences). Band intensities were assessed using associated Image Studio software.

### siRNA transfection

All siRNA transfections were performed using Lipofectamine RNAiMAX transfection reagent (Invitrogen), in accordance with the manufacturer’s protocol. Cells were seeded the day before treatment and 40 nM of siRNA was transfected. Cells were incubated for either 24, 48 or 72 h before data collection. Sequences of siRNAs used in this paper were as follows: ONTARGETplus Non-targeting siRNA oligo#1 (NT1) 5’-TGGTTTACATGTCGACTAA-3’ (Dharmacon, D-001810-01) and FlexiTube siRNA h USP31 oligo #1 (Q1) 5’-CCGAGTTCATGAAGACCTCAA-3’ (Qiagen, SI00758422), oligo #2 (Q2) 5’-CCCGAAATATTTAGGCCTGAA-3’ (Qiagen, SI00758415) and oligo 5 (Q4) 5’-CACCGAGCTCTTCGCCGAGTA-3’ (Qiagen, SI03058272).

### Plasmid DNA transfection

All plasmid DNA transfections were performed up to 24 h prior to collection of experimental data with cells at 60-80% confluency at the time of transfection. 1 μg of plasmid DNA (per six-well plate or ibidi dish) was transfected using Genejuice (Novagen) according to the manufacturer’s instructions. Cells were imaged the following day.

### Cell synchronisation

Cells were seeded into 10cm dishes and incubated for 2 days until 80% confluent before initiating synchronisation. Cells were synchronised using the following two protocols. i) For collection of cells in G1/S, S or G2 phase, cells were synchronised via a double thymidine block. First, 2 mM thymidine (Sigma, T1895) was added to the cell culture medium and cells incubated for 16 h. This first thymidine block was released for 8 h by exchanging for fresh medium before adding a second dose of 2mM thymidine for a further 16 h. Cells were released again into fresh medium and collected and lysed in urea lysis buffer at T=0 (G1), T=2.5 h (G1/Early S), T=5.5 h (Late S) and T=7.5 h (G2). ii) For collection of cells in mitosis, cells were treated with 2 mM thymidine for 24 h followed by release from thymidine into fresh medium containing 100 ng/ml nocodazole (Sigma, M1404) for 18 h. Mitotic cells were collected by pipetting the mitotic cells from the dish (“mitotic shake-off”). Cells were pelleted by centrifugation at 200 g for 5 min before resuspension in pre-warmed fresh medium to release from nocodazole. The cells were pelleted and washed a further 2x in fresh medium before a final resuspension in DMEM containing 25 mM HEPES pH 7.2. Samples were either lysed immediately or incubated for specified time points in a water bath at 37°C before being lysed on ice using urea lysis buffer. For experiments where USP31 depletion was performed, sychronisation was started 6 h after siRNA transfection.

### Lambda phosphatase

Samples were lysed in 2% SDS lysis buffer (2% SDS, 1mM EDTA, 50mM NaF) and sonicated before being subjected to a buffer exchange into lambda phosphatase buffer (150 mM NaCl, 1 mM EDTA, 1 mM EGTA, 1% Triton X-100, 20 mM Tris-HCl pH 7.5) using an Amicon^®^ Ultra 0.5 ml Centrifugal Filter following the manufacturers instructions. Protein dephosphorylation was achieved by incubation of 20 μg sample with 400 units of λ phosphatase (New England Biolabs) in 10X NEBuffer for protein MetalloPhosphatases (PMP) and 10mM MnCl2 in 50 μL at 30°C for 30min, prior to inactivation with SDS sample buffer.

### Immunofluorescence imaging

U2OS cells were grown on coverslips and treated as described for each experimental setup before being fixed with ice cold methanol at −20 °C for 5 min. Cells were then rinsed with PBS and non-specific sites were blocked using 10% goat serum in PBS for 30 min at room temperature (RT). Primary antibodies (1 h) and secondary antibodies (30 min) were applied sequentially at RT. Each antibody incubation was followed by 2 x 4 min washes with PBS. After a final wash in water, coverslips were mounted on glass slides using Mowiol supplemented with DAPI. Images were acquired using a Marianas spinning disk confocal microscope (3i, Intelligent Imaging Innovations, Germany) with either a Plan-Apochromat 40x/1.3NA Oil Objective or a Plan-Apochromat 63x/1.4NA Oil Objective. For immunofluorescence of the CPC localisation, 3D wide-field images were acquired using an AxioImager Z1 (100x Plan-Apochromatic oil differential interference contrast objective lens, 1.46 NA, ZEISS) equipped with a CCD camera (ORCA-R2, Hamamatsu) operated by Zen software (ZEISS) or a 3i Mariana spinning disk confocal microscope with a Plan-Apochromat 63x/1.4NA Oil Objective and a FLASH4 sCMOS Hamamatsu camera.

### PA tubulin

Kinetochore microtubule dynamics were determined by following the fluorescence decay of tubulin in U2OS cells expressing PA-GFP-α-tubulin/mCherry-tubulin. Cells were cultured in ibidi dishes, and USP31 was depleted using siRNA for 24 h prior to imaging. Cells were arrested in metaphase using 10 μM MG132 for 1 h prior to imaging, not exceeding 2 h by the end of imaging. Cells were imaged at 37 °C using a Plan-Apo 100× 1.4 NA DIC objective on a Nikon TE2000U inverted microscope equipped with a Yokogawa CSU-X1 spinning-disc confocal head containing two laser lines (488 nm and 561 nm) and a Mosaic (Andor) photoactivation system (405 nm). Briefly, metaphase cells were selected by imaging the brightfield and identifying the metaphase plate. Cherry-tubulin was used to confirm the mitotic spindle. A stripe was then generated in one half spindle, in proximity to the metaphase plate, and a z-stack of 7 planes, with a 1 μm step size was acquired for both the 488 and the 561 channels. The half-life of the microtubules was then calculated using a kymograph algorithm in MATLAB. Kymograph generation and analysis was as described in (Girao and Maiato, 2020) and (Ferreira et al., 2020). MT turnover was calculated based on a fitted curve of the normalized intensities at each time point (corrected for photobleaching) to a double exponential curve A1*exp(–k1*t) + A2*exp(–k2*t) using MATLAB (MathWorks), in which t is time, A1 represents the less stable (non-kMTs) population, and A2 the more stable (kMTs) population with decay rates of k1 and k2, respectively (cells displaying an R2 value <0.99 were excluded from quantification).

### Other Live cell imaging

All live cell imaging was performed in 35 mm Ibidi dishes (Ibidi, 81156) to monitor mitotic progression, U2OS mRFP-H2B cells were imaged using a Nikon Ti-Eclipse microscope and a CFI Super Plan Fluor ELWD ADM 20x/0.45NA objective (Fig. 2C).

Representative images of the spindle morphology in control and USP31 depleted U2OS cells (Fig. 4B) were captured using an Abberior ‘Expert Line’ gated-STED microscope, equipped with a Nikon Lambda Plan-Apo 1.4 NA 60x objective lens.

For imaging of misaligned and lagging chromosomes, GFP-H2B and Cherry-tubulin were selected at late Prometaphase and time-lapse image stacks of 2 μm steps were collected every minute at 37 °C using a temperature-controlled Nikon TE2000-E microscope equipped with a modified Yokogawa CSU-X1 spinning-disc head (Yokogawa Electric), an electron multiplying iXon+ DU-897 EM-CCD camera (Andor), a filter-wheel and an oil-immersion 100x 1.4 NA Plan-Apo DIC (Nikon). Two laser lines were used for excitation at 488 and 561nm. All image acquisition was controlled by NIS Elements AR software (Fig. 4E).

All other live cell imaging was performed using a Mariana 3i spinning disk confocal microscope and a FLASH4 sCMOS Hamamatsu camera. For imaging of mRFP-H2B expressing U2OS cells transiently transfected with GFP-USP31, time-lapse image stacks of 0.3 μm steps (40 slices) were collected every minute using a Plan-Apochromat 40x/1.3NA Oil Objective (Fig. 4A). Time-lapse image stacks (1 μm) of cells stably expressing wild-type and catalytically inactive GFP-USP31 under anaphase-like conditions in the presence of CDK1 inhibitor were collected every minute using a Plan Apochromat 100x/1.4NA Oil Objective (Fig. 5F). VSV-INCENP-GFP expressing U2OS cells were likewise imaged every minute (1 μm time-lapse steps) using a Plan-Apochromat 63x/1.4NA Oil Objective M27 (Fig. 7B).

### Flow cytometry

USP31 was knocked down in U2OS cells using four individual siRNAs for 48 h. The cell cycle was then profiled in these cells using flow cytometry. Briefly, the cells were trypsinised, pelleted and resuspended in ice cold ethanol. The fixed pellet was washed twice with PBS and stained with Propidium Iodine, and analysed on a ATTUNE Flow cytometer in conjunction with FlowJo software.

## Supplemental material

**Fig. S1** shows the localisation of USP31 to microtubules, centrosomes and cilia and maps the microtubule localisation determinants to its Ser-rich C-terminal region. It also provides evidence for widespread expression of USP31 in cell lines and for the depletion efficiency of the siRNA oligos deployed in this study. **Fig. S2** shows the mitotic phenotype analysis of U2OS cell lines stably overexpressing either wild-type or catalytically inactive, N-terminally GFP-tagged USP31 (C137A). **Fig. S3** provides additional evdidence for CDK1-dependent phosphorylation of USP31. **Fig. S4** provides further evidence for the impact of USP31 depletion as well as expression of catalytically inactive USP31 on the stability and localisation of CPC components during mitosis. **Video 1** (relating to Fig.4A) shows the rapid recruitment of GFP-USP31 onto kinetochore microtubules and the central spindle upon anaphase onset in U2OS cells stably expressing RFP-H2B. **Video 2** (relating to Fig. 5F) shows the rapid recruitment of GFP-USP31 to the central spindle upon CDK1 inhibition in U2OS cells arrested at metaphase. **Video 3** (relating to Fig. 5F) shows the rapid recruitment of catalytically inactive GFP-USP31 to the central spindle upon CDK1 inhibition followed by the appearance of multiple ectopic cleavage furrow sites. **Video 4 (**relating to Fig. 7B) shows the transition of Inducibly expressed INCENP-GFP from the kinetochores to the midzone upon anaphase onset in U2OS cells transfected with a control siRNA (NT1). **Video 5** and **6 (**relating to Fig. 7B) illustrate the delayed transition of inducibly expressed INCENP-GFP from kinetochores to the midzone in USP31 depleted cells transfected with siRNA Q1 and Q4, respectively.

## Acknowledgements

EB and JG have been funded by North West Cancer Research (CR1151 and CD2019.07) and HG by the Wellcome Trust (102172/B/13/Z). EB was the recipient of an EMBO Short-Term Fellowship 8457. MC is the recipient of a Royal Society Industry Fellowship (INF\R2\212031). We thank members of the Maiato Lab António Pereira, Hugo Girao, Bernardo Orr and Ana Almeida for help with microscopy and associated analysis.

Work in the Maiato lab is funded by an European Research Council (ERC) consolidator grant CODECHECK, under the European Union’s Horizon 2020 research and innovation programme (grant agreement 681443), Fundação para a Ciência e a Tecnologia of Portugal (PTDC/MED-ONC/3479/2020), and a La Caixa Health Research Grant (LCF/PR/HR21/52410025). We thank Dan Rigden (University of Liverpool) for assistance with sequence motif analysis and Susanne Lens (UMC Utrecht, the Netherlands) for generously providing the VSV-INCENP-GFP cell line.

## References

Adriaans, I.E., P.J. Hooikaas, A. Aher, M.J.M. Vromans, R.M. van Es, I. Grigoriev, A. Akhmanova, and S.M.A. Lens. 2020. MKLP2 Is a Motile Kinesin that Transports the Chromosomal Passenger Complex during Anaphase. Curr Biol. 30:2628–2637 e2629.

Barisic, M., P. Aguiar, S. Geley, and H. Maiato. 2014. Kinetochore motors drive congression of peripheral polar chromosomes by overcoming random arm-ejection forces. Nat Cell Biol. 16:1249–1256.

Carmena, M., M. Wheelock, H. Funabiki, and W.C. Earnshaw. 2012. The chromosomal passenger complex (CPC): from easy rider to the godfather of mitosis. Nat Rev Mol Cell Biol. 13:789–803.

Clague, M.J., I. Barsukov, J.M. Coulson, H. Liu, D.J. Rigden, and S. Urbe. 2013. Deubiquitylases from genes to organism. Physiol Rev. 93:1289–1315.

Clague, M.J., S. Urbe, and D. Komander. 2019. Breaking the chains: deubiquitylating enzyme specificity begets function. Nat Rev Mol Cell Biol. 20:338–352.

Clancy, A., C. Heride, A. Pinto-Fernandez, H. Elcocks, A. Kallinos, K.J. Kayser-Bricker, W. Wang, V. Smith, S. Davis, S. Fessler, C. McKinnon, M. Katz, T. Hammonds, N.P. Jones, J. O’Connell, B. Follows, S. Mischke, J.A. Caravella, S. Ioannidis, C. Dinsmore, S. Kim, A. Behrens, D. Komander, B.M. Kessler, S. Urbe, and M.J. Clague. 2021. The deubiquitylase USP9X controls ribosomal stalling. J Cell Biol.220 (3): e202004211.

Douanne, T., G. Andre-Gregoire, A. Thys, K. Trillet, J. Gavard, and N. Bidere. 2019. CYLD Regulates Centriolar Satellites Proteostasis by Counteracting the E3 Ligase MIB1. Cell Rep. 27:1657–1665 e1654.

Faesen, A.C., M.P.A. Luna-Vargas, P.P. Geurink, M. Clerici, R. Merkx, W.J. van Dijk, D.S. Hameed, F. El Oualid, H. Ovaa, and T.K. Sixma. 2011. The Differential Modulation of USP Activity by Internal Regulatory Domains, Interactors and Eight Ubiquitin Chain Types. Chemisty and Biology. 18:1550–1561.

Ferreira, L.T., A.C. Figueiredo, B. Orr, D. Lopes, and H. Maiato. 2018. Dissecting the role of the tubulin code in mitosis. Methods Cell Biol. 144:33–74.

Ferreira, L.T., B. Orr, G. Rajendraprasad, A.J. Pereira, C. Lemos, J.T. Lima, C. Guasch Boldu, J.G. Ferreira, M. Barisic, and H. Maiato. 2020. alpha-Tubulin detyrosination impairs mitotic error correction by suppressing MCAK centromeric activity. J Cell Biol. 219, (4):e201910064.

Gao, J., L. Huo, X. Sun, M. Liu, D. Li, J.T. Dong, and J. Zhou. 2008. The tumor suppressor CYLD regulates microtubule dynamics and plays a role in cell migration. J Biol Chem. 283:8802–8809.

Gavory, G., C.R. O’Dowd, M.D. Helm, J. Flasz, E. Arkoudis, A. Dossang, C. Hughes, E. Cassidy, K. McClelland, E. Odrzywol, N. Page, O. Barker, H. Miel, and T. Harrison. 2018. Discovery and characterization of highly potent and selective allosteric USP7 inhibitors. Nat Chem Biol. 14:118–125.

Girao, H., and H. Maiato. 2020. Measurement of Microtubule Half-Life and Poleward Flux in the Mitotic Spindle by Photoactivation of Fluorescent Tubulin. Methods Mol Biol.2101:235–246.

Gruneberg, U., R. Neef, R. Honda, E.A. Nigg, and F.A. Barr. 2004. Relocation of Aurora B from centromeres to the central spindle at the metaphase to anaphase transition requires MKlp2. J Cell Biol. 166:167–172.

Han, K.J., Z. Wu, C.G. Pearson, J. Peng, K. Song, and C.W. Liu. 2019. Deubiquitylase USP9X maintains centriolar satellite integrity by stabilizing pericentriolar material 1 protein. J Cell Sci. 132. (2):jcs221663

Heride, C., D.J. Rigden, E. Bertsoulaki, D. Cucchi, E. De Smaele, M.J. Clague, and S. Urbe. 2016. The centrosomal deubiquitylase USP21 regulates Gli1 transcriptional activity and stability. J Cell Sci. 129:4001–4013.

Honnappa, S., S.M. Gouveia, A. Weisbrich, F.F. Damberger, N.S. Bhavesh, H. Jawhari, I. Grigoriev, F.J. van Rijssel, R.M. Buey, A. Lawera, I. Jelesarov, F.K. Winkler, K. Wuthrich, A. Akhmanova, and M.O. Steinmetz. 2009. An EB1-binding motif acts as a microtubule tip localization signal. Cell. 138:366–376.

Janke, C., and G. Montagnac. 2017. Causes and Consequences of Microtubule Acetylation. Curr Biol. 27:R1287–R1292.

Kategaya, L., P. Di Lello, L. Rouge, R. Pastor, K.R. Clark, J. Drummond, T. Kleinheinz, E. Lin, J.P. Upton, S. Prakash, J. Heideker, M. McCleland, M.S. Ritorto, D.R. Alessi, M. Trost, T.W. Bainbridge, M.C.M. Kwok, T.P. Ma, Z. Stiffler, B. Brasher, Y. Tang, P. Jaishankar, B.R. Hearn, A.R. Renslo, M.R. Arkin, F. Cohen, K. Yu, F. Peale, F. Gnad, M.T. Chang, C. Klijn, E. Blackwood, S.E. Martin, W.F. Forrest, J.A. Ernst, C. Ndubaku, X. Wang, M.H. Beresini, V. Tsui, C. Schwerdtfeger, R.A. Blake, J. Murray, T. Maurer, and I.E. Wertz. 2017. USP7 small-molecule inhibitors interfere with ubiquitin binding. Nature. 550:534–538.

Kim, W., E.J. Bennett, E.L. Huttlin, A. Guo, J. Li, A. Possemato, M.E. Sowa, R. Rad, J. Rush, M.J. Comb, J.W. Harper, and S.P. Gygi. 2011. Systematic and Quantitative Assessment of the Ubiquitin-Modified Proteome. Mol Cell. 44:325–340.

Komander, D., F. Reyes-Turcu, L. J.D.F., P. Odenwaelder, K.D. Wilkinson, and D. Barford. 2009. Molecular discrimination of structurally equivalent Lys63-linked and linear polyubiquitin chains. EMBO Rep. 10:466–473.

Lamberto, I., X. Liu, H.S. Seo, N.J. Schauer, R.E. Iacob, W. Hu, D. Das, T. Mikhailova, E.L. Weisberg, J.R. Engen, K.C. Anderson, D. Chauhan, S. Dhe-Paganon, and S.J. Buhrlage. 2017. Structure-Guided Development of a Potent and Selective Non-covalent Active-Site Inhibitor of USP7. Cell Chem Biol. 24:1490–1500.

Li, J., V. D’Angiolella, E.S. Seeley, S. Kim, T. Kobayashi, W. Fu, E.I. Campos, M. Pagano, and B.D. Dynlacht. 2013. USP33 regulates centrosome biogenesis via deubiquitination of the centriolar protein CP110. Nature. 495:255–259.

Li, X., N. Song, L. Liu, X. Liu, X. Ding, X. Song, S. Yang, L. Shan, X. Zhou, D. Su, Y. Wang, Q. Zhang, C. Cao, S. Ma, N. Yu, F. Yang, Y. Wang, Z. Yao, Y. Shang, and L. Shi. 2017. USP9X regulates centrosome duplication and promotes breast carcinogenesis. Nat Commun. 8:14866.

Maerki, S., M.H. Olma, T. Staubli, P. Steigemann, D.W. Gerlich, M. Quadroni, I. Sumara, and M. Peter. 2009. The Cul3-KLHL21 E3 ubiquitin ligase targets aurora B to midzone microtubules in anaphase and is required for cytokinesis. J Cell Biol.187:791–800.

Nishi, R., P. Wijnhoven, C. le Sage, J. Tjeertes, Y. Galanty, J.V. Forment, M.J. Clague, S. Urbe, and S.P. Jackson. 2014. Systematic characterization of deubiquitylating enzymes for roles in maintaining genome integrity. Nature Cell Biology. 16:1016–1026.

Ramadan, K., R. Bruderer, F.M. Spiga, O. Popp, T. Baur, M. Gotta, and H.H. Meyer. 2007. Cdc48/p97 promotes reformation of the nucleus by extracting the kinase Aurora B from chromatin. Nature. 450:1258–1262.

Rusilowicz-Jones, E.V., F.G. Barone, F.M. Lopes, E. Stephen, H. Mortiboys, S. Urbe, and M.J. Clague. 2022. Benchmarking a highly selective USP30 inhibitor for enhancement of mitophagy and pexophagy. Life Sci Alliance. 5. (2) e202101287.

Song, L., A. Craney, and M. Rape. 2014. Microtubule-dependent regulation of mitotic protein degradation. Mol Cell. 53:179–192.

Sowa, M.E., E.J. Bennett, S.P. Gygi, and J.W. Harper. 2009. Defining the human deubiquitinating enzyme interaction landscape. Cell. 138:389–403.

Starita, L.M., Y. Machida, S. Sankaran, J.E. Elias, K. Griffin, B.P. Schlegel, S.P. Gygi, and J.D. Parvin. 2004. BRCA1-dependent ubiquitination of gamma-tubulin regulates centrosome number. Mol Cell Biol. 24:8457–8466.

Sumara, I., M. Quadroni, C. Frei, M.H. Olma, G. Sumara, R. Ricci, and M. Peter. 2007. A Cul3-based E3 ligase removes Aurora B from mitotic chromosomes, regulating mitotic progression and completion of cytokinesis in human cells. Dev Cell. 12:887–900.

Takahashi, H., S. Yamanaka, S. Kuwada, K. Higaki, K. Kido, Y. Sato, S. Fukai, F. Tokunaga, and T. Sawasaki. 2020. A Human DUB Protein Array for Clarification of Linkage Specificity of Polyubiquitin Chain and Application to Evaluation of Its Inhibitors. Biomedicines. 8(6):152.

Turnbull, A.P., S. Ioannidis, W.W. Krajewski, A. Pinto-Fernandez, C. Heride, A.C.L. Martin, L.M. Tonkin, E.C. Townsend, S.M. Buker, D.R. Lancia, J.A. Caravella, A.V. Toms, T.M. Charlton, J. Lahdenranta, E. Wilker, B.C. Follows, N.J. Evans, L. Stead, C. Alli, V.V. Zarayskiy, A.C. Talbot, A.J. Buckmelter, M. Wang, C.L. McKinnon, F. Saab, J.F. McGouran, H. Century, M. Gersch, M.S. Pittman, C.G. Marshall, T.M. Raynham, M. Simcox, L.M.D. Stewart, S.B. McLoughlin, J.A. Escobedo, K.W. Bair, C.J. Dinsmore, T.R. Hammonds, S. Kim, S. Urbe, M.J. Clague, B.M. Kessler, and D. Komander. 2017. Molecular basis of USP7 inhibition by selective small-molecule inhibitors. Nature. 550:481–486.

Urbe, S., H. Liu, S.D. Hayes, C. Heride, D.J. Rigden, and M.J. Clague. 2012. Systematic survey of deubiquitinase localisation identifies USP21 as a regulator of centrosome and microtubule associated functions. Mol Biol Cell. 23:1095–1103.

van der Waal, M.S., A.T. Saurin, M.J. Vromans, M. Vleugel, C. Wurzenberger, D.W. Gerlich, R.H. Medema, G.J. Kops, and S.M. Lens. 2012. Mps1 promotes rapid centromere accumulation of Aurora B. EMBO Rep. 13:847–854.

Vong, Q.P., K. Cao, H.Y. Li, P.A. Iglesias, and Y. Zheng. 2005. Chromosome alignment and segregation regulated by ubiquitination of survivin. Science. 310:1499–1504.

Xu, G., J.S. Paige, and S.R. Jaffrey. 2010. Global analysis of lysine ubiquitination by ubiquitin remnant immunoaffinity profiling. Nat Biotechnol. 28:868–873.

Ye, Y., H. Scheel, K. Hofmann, and D. Komander. 2009. Dissection of USP catalytic domains reveals five common insertion points. Mol Biosyst. 5:1797–1808.

