## Supplementary Material for "The deubiquitylase USP31 controls the Chromosomal Passenger Complex and spindle dynamics"

**Supplemental Figure legends**

**Figure S1: USP31 localises to microtubules, centrosomes and primary cilia**

**A.** hTERT-RPE1 (left) or NIH3T3 cells (right) were transfected with GFP-USP31, fixed with ice-cold MeOH and stained as indicated for Pericentrin (centrosomes) or Arl13b (primary cilia). NIH3T3 cells were serum-starved for 24 h prior to fixation to induce cilia formation. Images were captured with a 3i spinning disk confocal microscope. Scale bar: 10 $\mu$m.

**B and C.** Localisation and relative expression of USP31 deletion mutants depicted in Figure 1C. U2OS cells were transfected with GFP-tagged USP31 deletion mutants, fixed with ice-cold MeOH and stained for α-tubulin. Images were captured with a 3i spinning disk confocal microscope. Scale bar: 10 $\mu$m. C. Lysates of cells shown in B were analysed by western blotting.

**D:** Total lysates from the indicated cell lines were analysed by western blotting and probed for USP31.

**E.** Western blot showing the knockdown efficiency for U2OS cells treated for 72 h with siRNA targeting USP31 (Q1, Q2, Q4) or non-targeting siRNA (NT1).

**Figure S2: Immunofluorescence microscopy of stable cell lines overexpressing either wild-type or catalytically inactive USP31.**

**A.** U2OS cells stably over-expressing either wild type (WT8, WT9, WT13) or catalytically inactive (CA1, CA7) GFP-USP31 were fixed with MeOH and stained for α-tubulin. Images were acquired using a 3i spinning disk confocal. Scale bar: 10 $\mu$m

**B.** U2OS cells stably expressing either wild type (WT8, WT9, WT13) or catalytic inactive (CA1, CA7) GFP-USP31 were fixed with MeOH and stained for γ-tubulin (red) and Pericentrin (blue). Images were acquired using a 3i spinning disk confocal. Scale bar: 10 $\mu$m.

**Figure S3: USP31 phosphorylation is dependent on CDK1 activity.**

**A.** USP31 phosphorylation occurs as cells enter mitosis. U2OS cells were treated with thymidine for 24 h before being released into fresh DMEM containing nocodazole and incubated for the indicated time points.

**B.** USP31 dephosphorylation and decay occurs as cells exit mitosis. U2OS cells were synchronised with a single thymidine block followed by nocodazole to prometaphase and then released into fresh DMEM to allow cells to proceed through mitosis.

**C.** U2OS cells were synchronised as in B, then released from nocodazole into fresh DMEM. Samples were treated with either the Aurora B inhibitor ZM447439 (10 $\mu$M) or the CDK1 inhibitor RO3306 (10 $\mu$M), or DMSO for the indicated timepoints.

**D.** U2OS cells were synchronised to prometaphase via single thymidine block followed by nocodazole and released into fresh DMEM. Samples were treated immediately with either DMSO, RO3306 (RO, 10 $\mu$M) or ZM447439 (ZM, 10 $\mu$M) for 30 min, or were first pre-treated with MG132 (MG, 5$\mu$M) for 1 h before RO or ZM addition.

**E.** Schematic diagram showing the incubation schedule for samples shown in F and Figures 5G.

**F.** U2OS cells were synchronised using a single thymidine block followed by nocodazole and released into fresh DMEM. One sample was lysed immediately (prometa) whereas MG132 (MG, 5 µM) was added to all other samples and incubated for specified times. For control cells, MG132 washout in fresh DMEM was performed and cells incubated for specified times to allow re-entry into mitosis. Asy: Asynchronous cell lysates. Blue arrows represent full length USP31 whereas green arrows represent a proposed shorter isoform. Top arrows (light) and bottom arrows (dark) indicate the phosphorylated and unphosporylated USP31 species respectively.

**Figure S4: USP31 modulates CPC stabilisation and localisation during mitosis**

**A.** U2OS cells were transfected with siRNA against USP31 (Q1) or a non-targeting control (NT1) for 48 h, then treated with MG132 (5 $\mu$M) for 1 h and subsequently treated with RO3306 (10 $\mu$M) for 8 min. Cells were fixed with MeOH and stained for the indicated antibodies. Images acquired with a 3i spinning disk confocal microscope, 63x objective. Single slices from representative images are shown. Scale bar 10 $\mu$M.

**B.** U2OS cells were transfected with siRNA against USP31 (Q1 or Q4) or a non-targeting control (NT1) for 48 h, and synchronised to prometaphase with a single thymidine block followed by nocodazole. Cells were then washed and released into fresh DMEM for 2-3 hours to allow progression to anaphase. Cells were fixed with

MeOH and stained for the indicated proteins. Images were acquired with a 3i spinning disk confocal. Single slices from representative images are shown. Scale bar = 10 $\mu$m.

**C.** Parental U2OS cells (Par) or cells stably expressing wild-type (WT13) or catalytic inactive (CA1) USP31 were synchronised at prometaphase (Prometa) using thymidine and nocodazole and lysed alongside asynchronously grown cells (Asy). Samples were analysed by western blotting and probed for indicated proteins.

**D.** Quantification of western blots illustrated in C. Graph shows results from 3 independent experiments. Statistical analysis carried out via one-way ANOVA with a Dunnett's multiple comparisons test. ns not significant, *p≤0.05, **p≤0.01, ***p≤0.001, ****p≤0.0001.

**Supplementary Video legends**

**Video 1**
**USP31 localises to the mitotic spindle**
U2OS cells stably expressing mRFP-H2B (not shown) were transfected with GFP-USP31 for 21 h. Z-stacks (40 slices, 0.3 µm steps) were acquired every minute using a 3i-spinning disc confocal microscope. Time is in hours:minutes. Scale bar = 10 µm. A small amount of GFP-USP31 is seen localised at the metaphase spindle, with a rapid recruitment onto kinetochore microtubules and the central spindle apparent upon anaphase onset. Relates to Fig. 4A.

**Video 2**
**Localisation of USP31 at metaphase is controlled by CDK1-dependent phosphorylation**
U2OS cells stably expressing GFP-tagged wild-type USP31 (WT13) were arrested in metaphase using MG132 (5 µM) for 1 h. Cells were immediately imaged following the addition of RO3306 (CDK1 inhibitor; 10 µM) using a 3i-spinning disc confocal microscope. Z-stacks (15 slices, 1 µm steps) were acquired every minute. Time is in hours:minutes. Scale bar = 10 µm. GFP-USP31 is rapidly recruited to the central spindle upon CDK1 inhibition. Relates to Fig. 5F.

**Video 3**
**Localisation of USP31 at metaphase is controlled by CDK1-dependent phosphorylation and causes multiple ectopic furrowing in U2OS cells stably expressing GFP-tagged catalytically inactive USP31 (CA1).** Cells were arrested in metaphase using MG132 (5 µM) for 1 h then immediately imaged following the addition of RO3306 (CDK1 inhibitor; 10 µM) using a 3i-spinning disc confocal microscope. Z-stacks (15 slices, 1 µm steps) were acquired every minute. Time is in hours:minutes. Scale bar = 10 µm. Catalytically inactive GFP-USP31 is rapidly recruited to the central spindle upon CDK1 inhibition and multiple ectopic furrowing sites are observed. Relates to Fig. 5F.

**Video 4**
**USP31 depletion destabilises ectopically expressed INCENP-GFP and delays its translocation to the midzone.** U2OS cells inducibly expressing INCENP-GFP were transfected with siRNA against a non-targeting control (NT1) for 45 h, then treated with doxycycline (1 µg/ml) for 3 h to induce INCENP-GFP expression. Cells were imaged using a 3i-spinning disc confocal microscope. Z-stacks (18 slices, 1 µm steps) were acquired every minute. Time is in hours:minutes. Scale bar = 10 µm. Inducibly expressed INCENP-GFP transitions from the kinetochores to the midzone and then the midbody upon anaphase onset in control cells. Relates to Fig 7B, showing frames 5-16.

**Video 5**
**USP31 depletion destabilises ectopically expressed INCENP-GFP and delays its translocation to the midzone.** U2OS cells inducibly expressing INCENP-GFP were transfected with siRNA against USP31 (Q1) for 45 h, then treated with doxycycline (1 µg/ml) for 3 h to induce INCENP-GFP expression. Cells were imaged using a 3i-spinning disc confocal microscope. Z-stacks (18 slices, 1 µm steps) were acquired every minute. Time is in hours:minutes. Scale bar = 10 µm. Transition of inducibly expressed INCENP-GFP from kinetochores to the midzone is delayed in USP31 depleted cells. Relates to Fig 7B, showing frames 5-16.

**Video 6**

**USP31 depletion destabilises ectopically expressed INCENP-GFP and delays its translocation to the midzone.** U2OS cells inducibly expressing INCENP-GFP were transfected with siRNA against USP31 (Q4) for 45 h, then treated with doxycycline (1 µg/ml) for 3 h to induce INCENP-GFP expression. Cells were imaged using a 3i-spinning disc confocal microscope. Z-stacks (18 slices, 1 µm steps) were acquired every minute. Time is in hours:minutes. Scale bar = 10 µm. Transition of inducibly expressed INCENP-GFP from kinetochores to the midzone is delayed in USP31 depleted cells. Relates to Fig 7B, showing frames 5-16.

### Figure S1

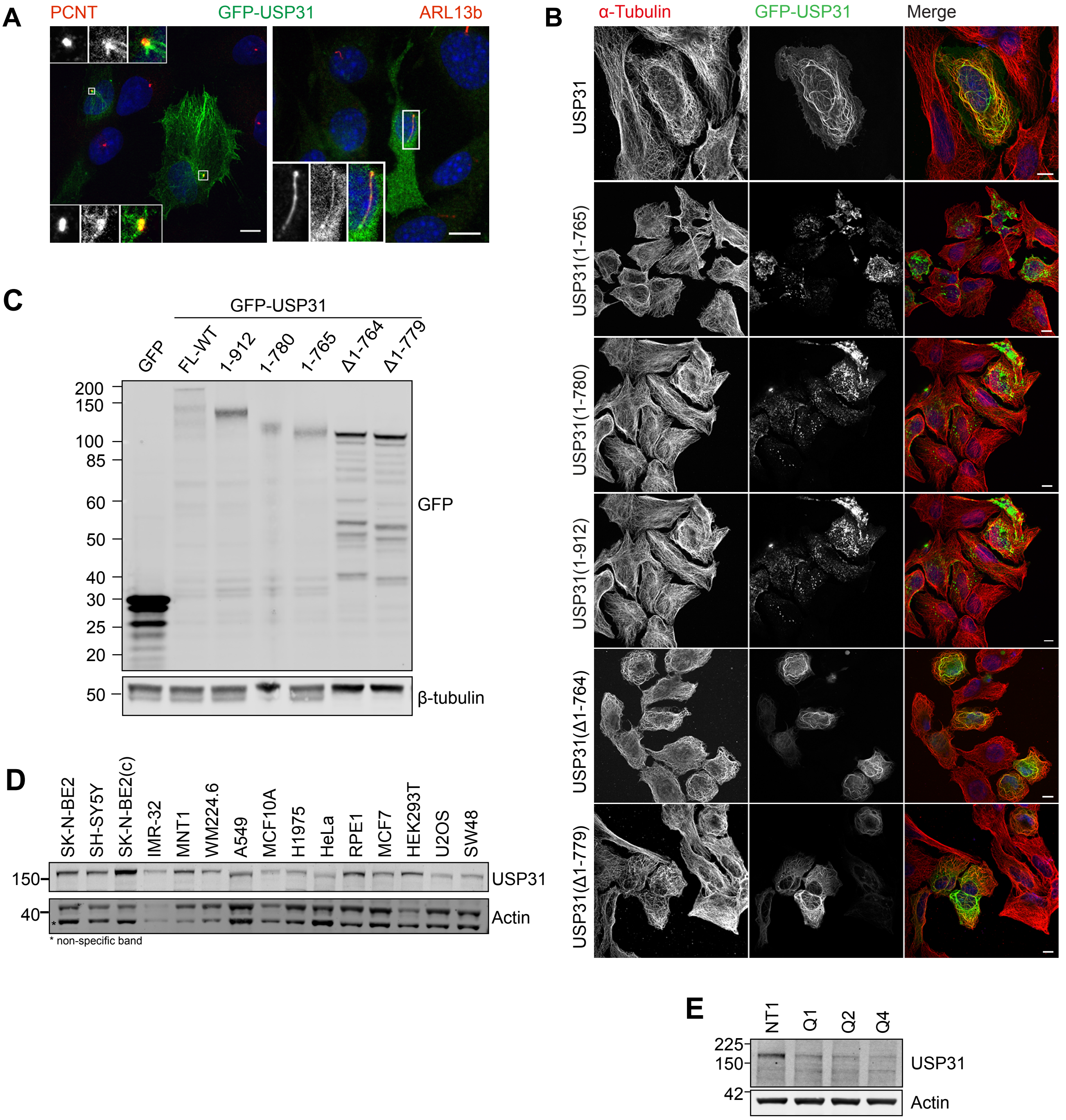
A
PCNT
GFP-USP31
ARL13b
B
α-Tubulin
GFP-USP31
Merge
USP31
USP31(1-765)
USP31(1-780)
USP31(1-912)
USP31(Δ1-764)
USP31(Δ1-779)
C
GFP-USP31
GFP
FL-WT
1-912
1-780
1-765
Δ1-764
Δ1-779
200
150
100
85
60
50
40
30
25
20
GFP
50
β-tubulin
D
SK-N-BE2
SH-SY5Y
SK-N-BE2(c)
IMR-32
MNT1
WM224.6
A549
MCF10A
H1975
HeLa
RPE1
MCF7
HEK293T
U2OS
SW48
150
USP31
40
Actin
* non-specific band
E
NT1
Q1
Q2
Q4
225
150
USP31
42
Actin

#### Figure S2

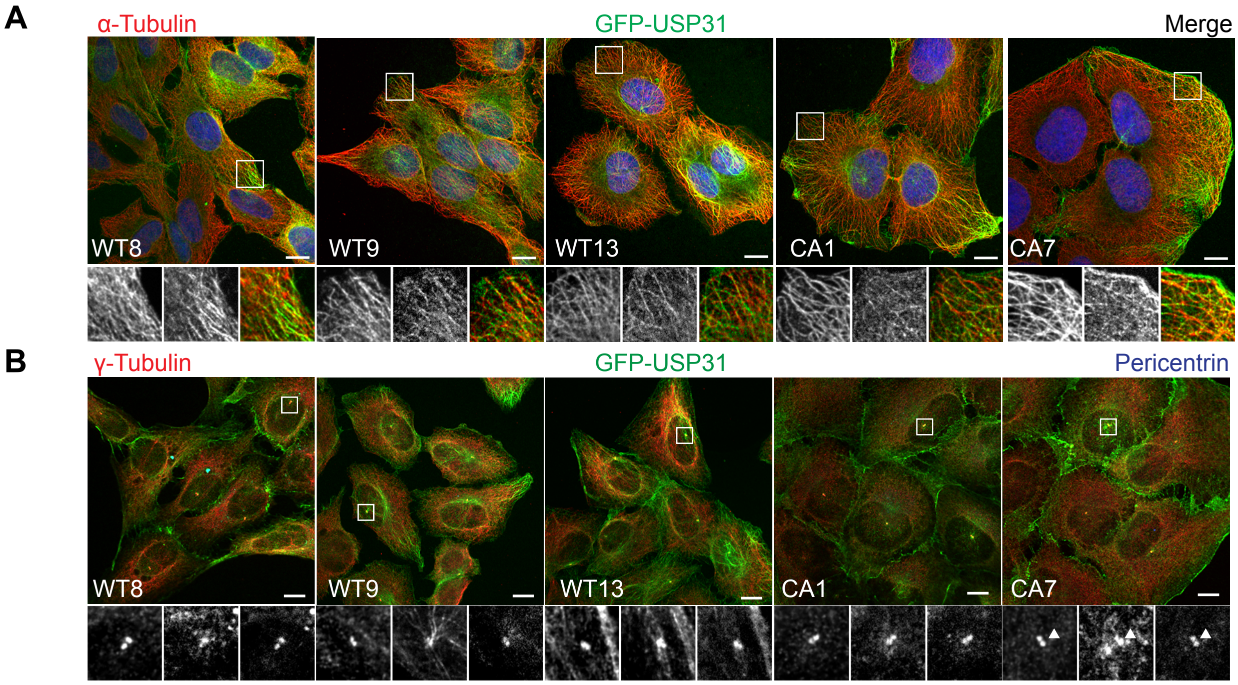

A
α-Tubulin
GFP-USP31
Merge
WT8
WT9
WT13
CA1
CA7
B
γ-Tubulin
GFP-USP31
Pericentrin
WT8
WT9
WT13
CA1
CA7

**Figure S3**

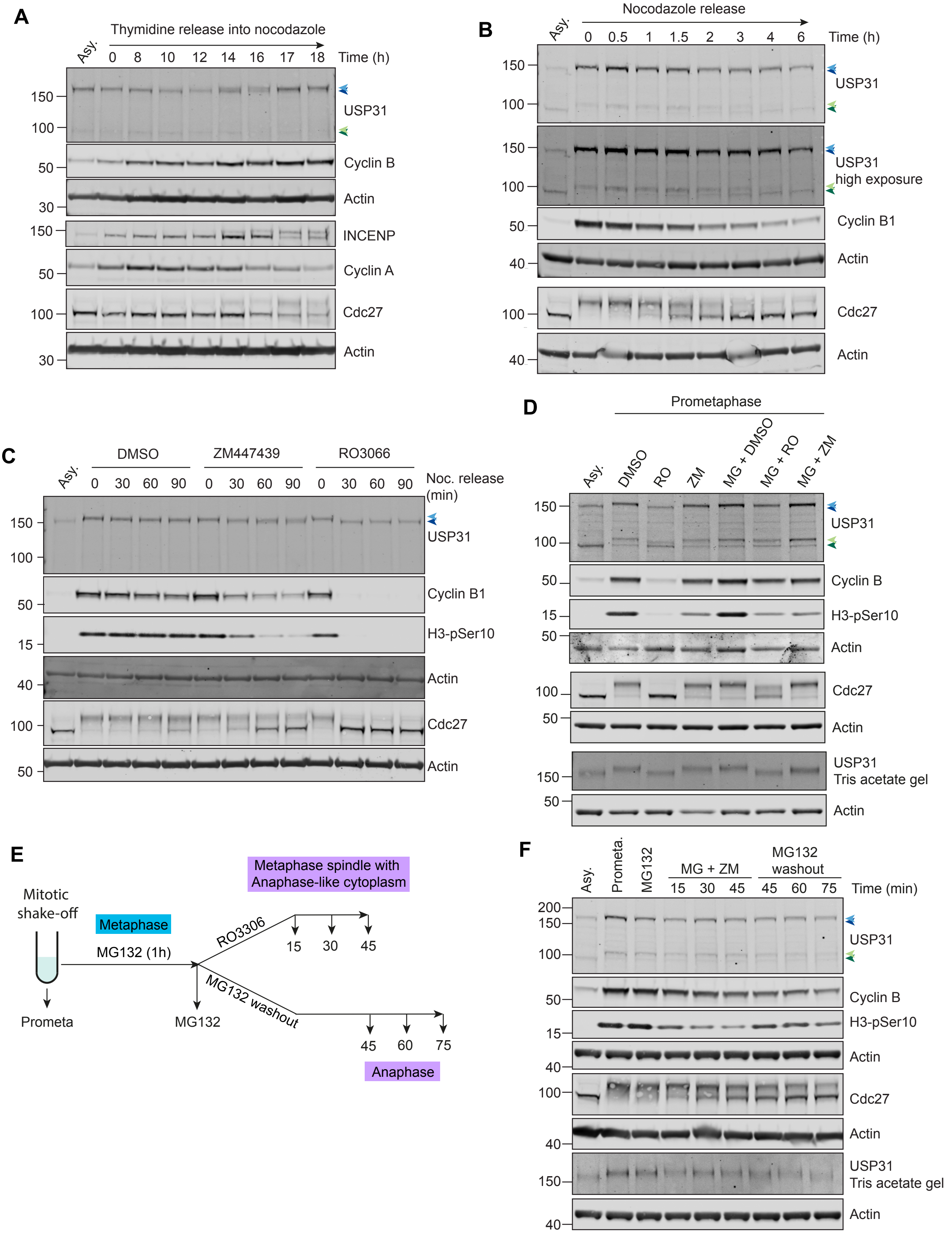
A
Thymidine release into nocodazole
Asy.
0 8 10 12 14 16 17 18 Time (h)
USP31
Cyclin B
Actin
INCENP
Cyclin A
Cdc27
Actin
B
Nocodazole release
0 0.5 1 1.5 2 3 4 6 Time (h)
USP31
USP31 high exposure
Cyclin B1
Actin
Cdc27
Actin
C
DMSO
ZM447439
RO3066
0 30 60 90
Noc. release (min)
USP31
Cyclin B1
H3-pSer10
Actin
Cdc27
Actin
D
Prometaphase
Asy.
DMSO
RO
ZM
MG + DMSO
MG + RO
MG + ZM
USP31
Cyclin B
H3-pSer10
Actin
Cdc27
Actin
USP31 Tris acetate gel
Actin
E
Mitotic shake-off
Metaphase
MG132 (1h)
Prometa
MG132
RO3306
15 30 45
MG132 washout
45 60 75
Metaphase spindle with Anaphase-like cytoplasm
Anaphase
F
Asy.
Prometa.
MG132
MG + ZM
MG132 washout
15 30 45
45 60 75
Time (min)
USP31
Cyclin B
H3-pSer10
Actin
Cdc27
Actin
USP31 Tris acetate gel
Actin

### Figure S4

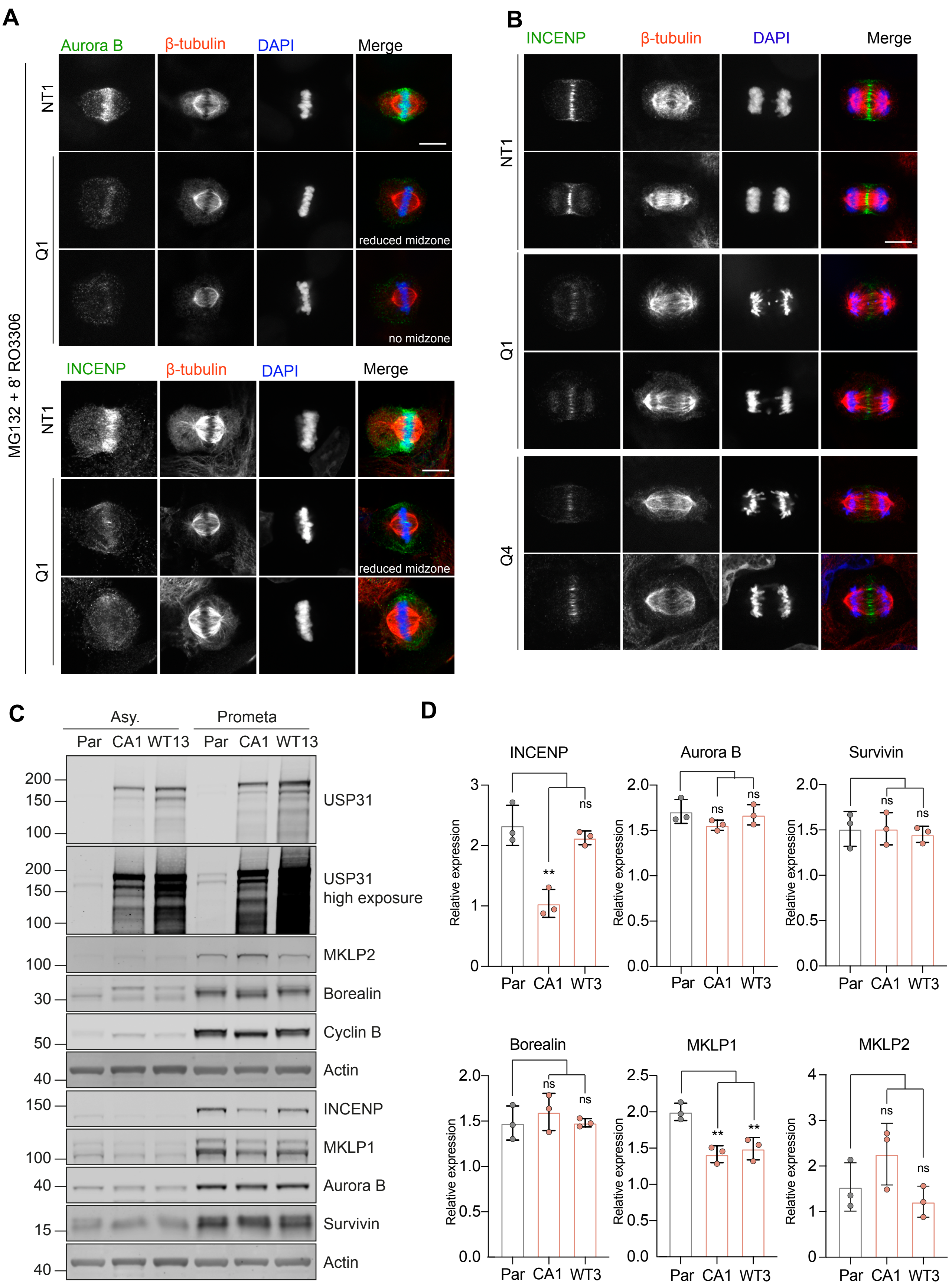

A
Aurora B
β-tubulin
DAPI
Merge
NT1
Q1
reduced midzone
no midzone
INCENP
MG132 + 8' RO3306
B
INCENP
β-tubulin
DAPI
Merge
NT1
Q1
Q4
C
Asy.
Prometa
Par CA1 WT13 Par CA1 WT13
USP31
USP31 high exposure
MKLP2
Borealin
Cyclin B
Actin
INCENP
MKLP1
Aurora B
Survivin
Actin
D
INCENP
Aurora B
Survivin
Borealin
MKLP1
MKLP2
Relative expression
Par CA1 WT3
ns
**
